# Ribosomes act as cryosensors in plants

**DOI:** 10.1101/2020.12.07.414789

**Authors:** David Guillaume-Schöpfer, Katja E. Jaeger, Feng Geng, Fabrizio G. Doccula, Alex Costa, Alex A. R. Webb, Philip A. Wigge

## Abstract

Cold temperatures are a threat to temperate plants, and *Arabidopsis thaliana* has acquired an adaptive gene expression network controlled by CBF transcription factors. The CBFs are sufficient to enable plants to survive otherwise lethal subzero temperatures. Constitutive CBF expression causes delayed flowering and stunted growth, and plants have evolved the ability to restrict CBF expression to occur only in the cold. This allows plants to anticipate likely freezing events and selectively deploy cold tolerance. The mechanism by which cold stress is sensed is however unknown. Here we show that protein translation rates in plants are proportional to temperature, and reduced translation rates trigger a rise in intracellular free calcium that activates the CAMTA transcription factors, and these directly activate cold-induced gene expression.

---

Freezing stress is a major threat to temperate plants, and *Arabidopsis thaliana* has evolved an adaptive transcriptional response activated by the C-REPEAT BINDING FACTOR1-3 (CBF1-3) AP2-type transcription factors, enabling survival of otherwise lethal subzero temperatures (Jaglo-Ottosen et al., 1998). Constitutive expression of CBFs however greatly reduces growth, and plants have evolved a pathway to restrict CBF expression to low temperatures (Jaglo-Ottosen et al., 1998). Activation of the CBFs by cold requires the activity of the *CALMODULIN-BINDING TRANSCRIPTION ACTIVATOR (CAMTA) 1, 2, 3* and *5* genes (Kim et al., 2013; Ohama et al., 2015). The CAMTAs are activated by Ca^2+^-calmodulins (Bouché et al., 2005), and calcium levels rise during cold stress (Plieth et al., 1999; Monroy et al., 1993; Knight et al., 1996; Berberich and Kusano, 1997; Tähtiharju et al., 1997; Yamazaki et al., 2008). The cryosensory mechanism in this pathway is however unknown. Here we show that ribosome translation rates provide the initial temperature sensing events necessary for the activation of the cold stress response.

## Translation rate is proportional to temperature in plants

Protein synthesis by ribosomes is a key cellular process, and the translation elongation rate is proportional to temperature in *E. coli* (Farewell and Neidhardt, 1998). To see if translation rates in plants are temperature-dependent, we measured the rate of protein synthesis using a cell-free system coupled to a Luciferase assay. This shows that translation efficiency increases strongly between 4 °C and 22 °C (Fig. 1A). We confirmed this effect by measuring protein levels (Supp. Fig. 1A).

**Figure 1:**
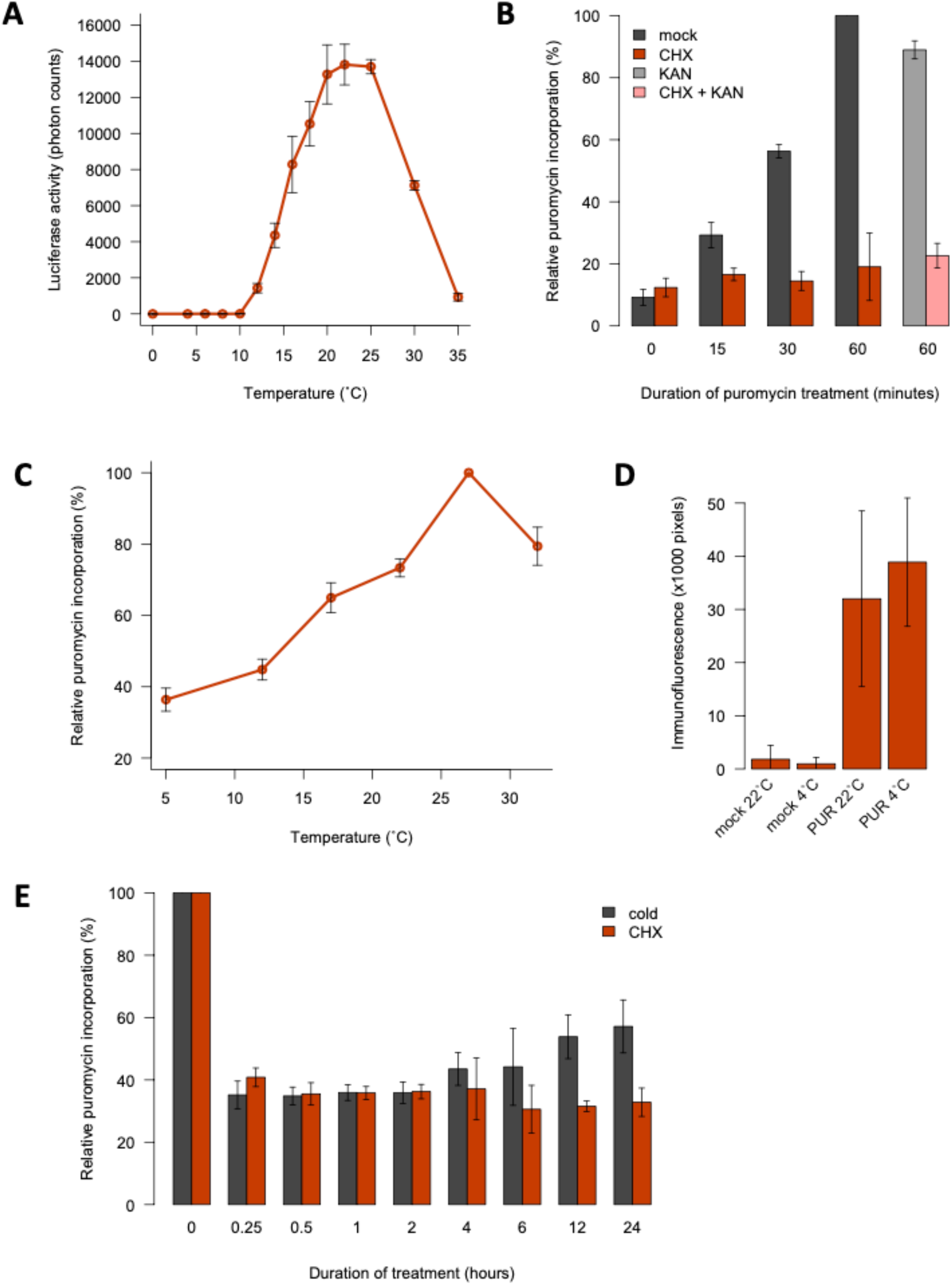
Translation rate is proportional to temperature. **A:** Luminescence from luciferase synthesised *in vitro* at a range of temperatures (4 replicates). **B, C, E:** Quantification of *in vivo* translation in Arabidopsis seedlings (?3 replicates, 10-15 seedlings each). **B:** Validation of the SUnSET assay. Seedlings were treated with 150 μM puromycin (PUR) following 60-minute incubations in CHX and/or kanamycin (KAN) at 30 μM or 0.1% DMSO (mock). **C:** Low temperatures reduce protein synthesis *in vivo*. Assays were performed five days after transferring 22°C-grown seedlings to the indicated temperatures. **D:** Low temperatures do not inhibit PUR uptake. Intracellular PUR was detected by immunofluorescence microscopy in seedlings at 4°C or 22°C (7 replicates). **E:** Cold and CHX cause rapid translational repression. Seedlings were treated with 0.1% DMSO at 22°C (mock) or 4°C or with 30 μM CHX at 22°C for 0 to 24 hours. PUR incorporation was normalised to mock controls. For all experiments, error bars indicate standard deviation.

To determine if temperature impacts translation globally in the cell, we assayed incorporation of the aminonucleoside puromycin in *de novo* synthesised peptides (EK et al., 2009). As expected, inhibiting 80S ribosomes sharply decreases translation (Fig. 1B). The results confirm that protein synthesis rates are proportional to temperature in plants (Fig. 1C). Temperature does not affect puromycin uptake (Fig. 1D, Supp. Fig. 1B), confirming the direct kinetic link between temperature and ribosome processivity in plants.

Translation declines rapidly in the cold, comparable to the effect of 30 μM CHX, consistent with the effect of temperature being direct (Fig. 1C,E). Translation rates begin to recover after 4 hours in the cold, indicating that acclimation mechanisms exist to enable some essential protein synthesis (Fig. 1C,E).

## Reducing ribosomal activity induces early cold stress signalling

Since protein synthesis rates are proportional to temperature, we tested if translation rates provide a mechanism to sense cold stress. Strikingly, slowing down translation using CHX is sufficient to rapidly induce the cold-responsive *pCBF2::LUC* reporter at room temperature, resembling the cold response (Fig. 2A,B), consistent with previous observations (Zarka et al., 2003; Berberich and Kusano, 1997). The response to CHX is specific, since later responding cold genes such as *COR15a* are not induced (Fig. 2B).

**Figure 2:**
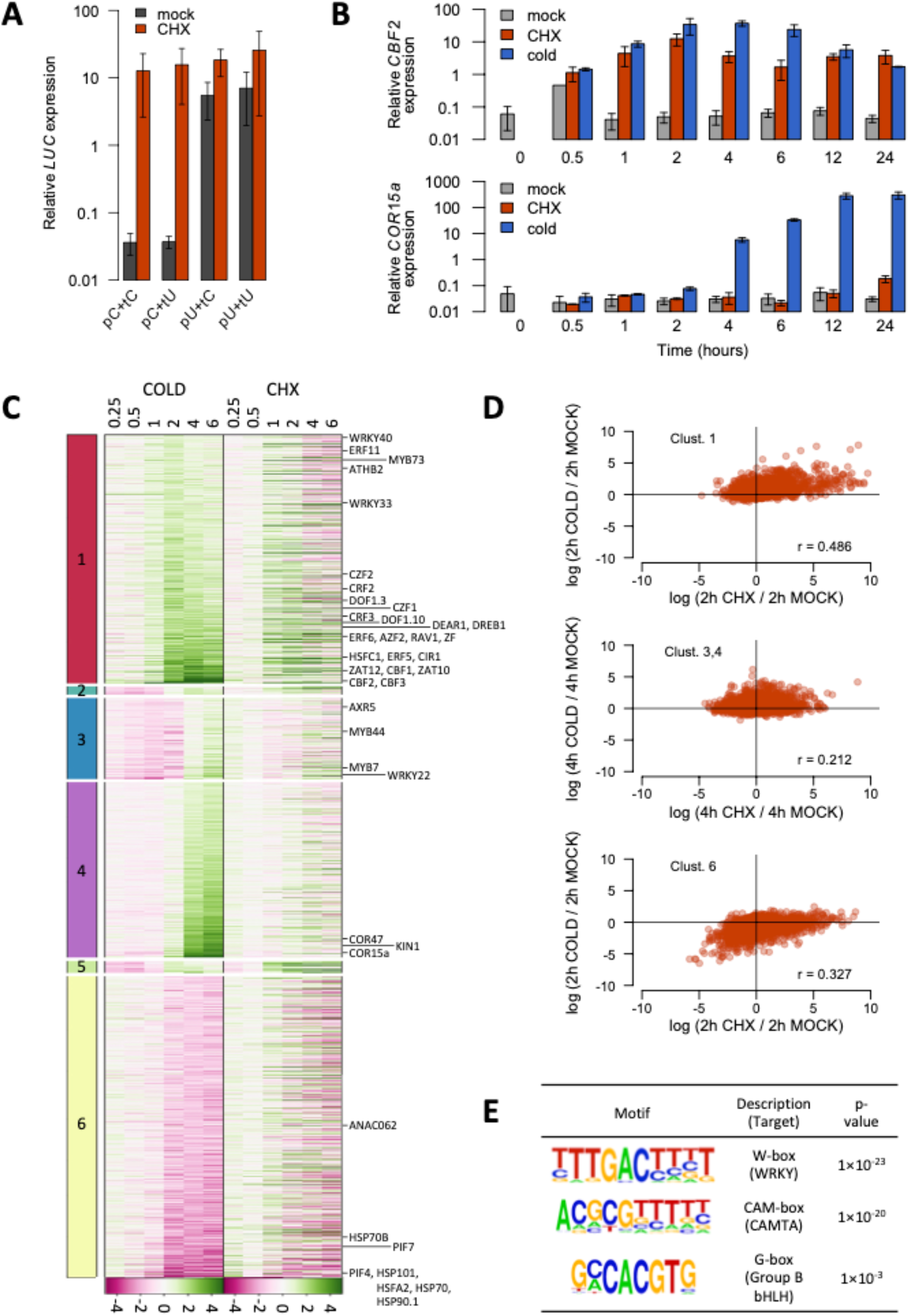
Cold signalling is activated by reduced ribosome activity. **A:** CHX activates the *CBF2* promoter. Expression of luciferase *(LUC)* driven from *CBF2* or *UBQ10* promoter (pC, pU) and terminator (tC, tU) sequences in Arabidopsis seedlings treated for 2 hours with CHX (30 μM, 22°C) or mock treatment (0.1% DMSO, 22°C) (four pools of 4-6 selected T1 seedlings). **B:** CHX induces *CBF2* but not *COR15a* expression. CHX, mock and cold (0.1% DMSO, 4°C) treatments were carried out for 0 to 24 hours (3 replicates, 10-15 seedlings each). Error bars indicate standard deviation (A, B). **C:** CHX induces an early cold-responsive cluster of genes. RNA-seq of seedlings after CHX, mock or cold treatments for 0.25 to 6 hours. Genes were hierarchically clustered according to their temporal pattern of cold induction. Colours indicate log2 fold-changes in expression relative to mock controls and genes of interest are annotated. **D:** Correlation between cold- and CHX-responsive transcriptomes for key clusters. Pearson correlation coefficients are indicated. **E:** Top three motifs enriched in promoters upregulated by 1-hour CHX treatment.

To determine if protein translation rates control the global response to cold stress, we compared the cold- and CHX-responsive transcriptomes over a 6 h time-course. Clustering of the cold-responsive genes reveals three major categories of response (Fig. 2C). Cluster 1 contains genes activated by 1-2 h of cold (early-inducible genes), including *CBF1*, *CBF2* and *CBF3*, and other transcription factors previously shown to respond rapidly to cold such as *ZAT10, ZAT12, ZF* and *HSFC1* (Park et al., 2015). Clusters 3 and 4 are activated at 4-6 h of cold and include *COR15a* and *COR47.* The majority of genes in Cluster 1 are rapidly induced by CHX treatment, whereas this is not the case for Clusters 3 and 4 (Fig. 2C,D). Cluster 6 contains genes that are repressed by cold, including many heat-inducible genes such as *HSP70*, *HSP101* and *HSFA2*, and many of these are repressed by CHX (Fig. 2C,D). CHX therefore activates the early cold-responsive transcriptional programme. Accordingly, genes induced after one hour of CHX treatment are enriched for promoter motifs targeted by transcription factors involved in cold signalling (Fig. 2E) (Doherty et al., 2009; Kidokoro et al., 2009; Lee and Thomashow, 2012; Kim et al., 2013; Park et al., 2015).

## Activation of cold- and CHX-responsive genes is mediated by CAMTAs

Cold-mediated *CBF* expression is dependent on the CAMTA family of transcription factors (Kim et al., 2013; Kidokoro et al., 2017), and we find that this is also the case for CHX (Fig. 3A). This suggests that CHX-induced *CBF* expression acts via the same pathway as cold stress. The CBF regulon is gated by the circadian clock, and we observe that *PSEUDO-RESPONSE REGULATOR 5, 7* and *9 (PRR5, 7* and *9)* regulate *CBF2* inducibility by CHX (Fig. 3B). *CBF2* expression is also more responsive to CHX at the beginning of the day, as has been described for the cold response, consistent with the CHX pathway acting through the same pathway as the cold stress signalling pathway (Supp. Fig. 2).

**Figure 3:**
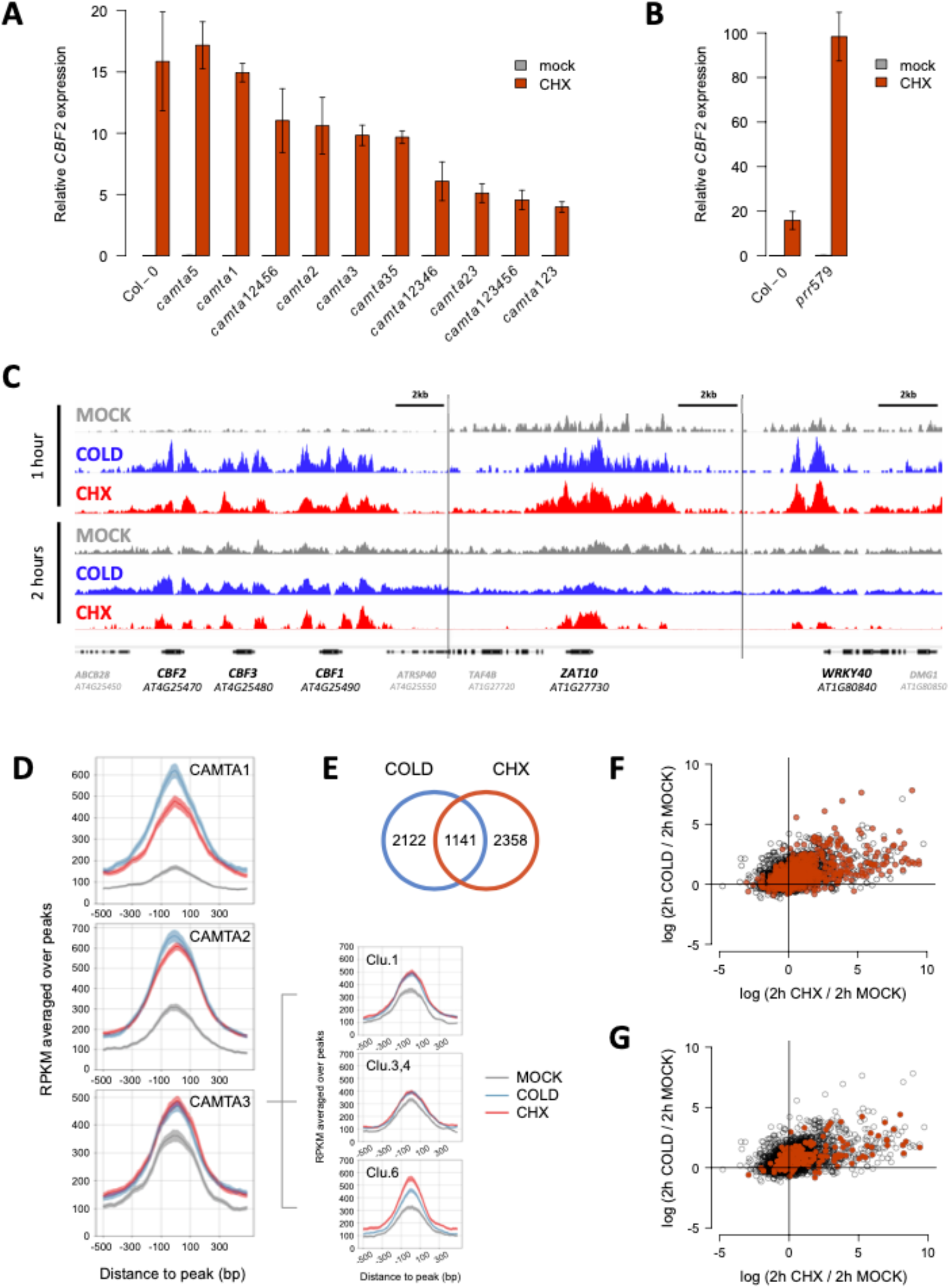
CAMTAs directly activate cold and CHX responsive genes. **A, B:** Arabidopsis *camta* (**A**) or *prr* mutants (**B**) were treated for 2 hours with CHX (30 μM, 22°C) or mock treatment (0.1% DMSO, 22°C). Error bars indicate standard deviation (3 replicates, 10-15 seedlings each). **C-G:** ChlP-seq of CAMTA1, 2 and 3 in complemented *camta123* seedlings after 1 or 2 hours of CHX, mock or cold (0.1% DMSO, 4°C) treatments. **C:** Cold- and CHX-associated binding of CAMTA2 at early cold-inducible genes *CBF1, CBF2, CBF3, ZAT10* and *WRKY40.* **D:** Genome-wide pile-ups of CAMTA1, 2 and 3 after 1-hour treatments, and cluster-specific pile-ups of CAMTA3. **E:** More than a third of genes bound by CAMTAs at 4°C are also bound during CHX treatments at 22°C. Numbers indicate genes at which there is an increase in occupancy of CAMTA1, 2 or 3 during 1 or 2 hours of treatments relative to mock controls. **F, G:** Many early cold-inducible genes are bound by CAMTAs. Log2 fold-changes in expression of genes from Cluster 1 during 2-hour CHX or cold treatments relative to mock treatments. Genes bound by CAMTA1, 2 or 3 in response to both cold and CHX (**F**) or during either treatment (**G**) are indicated in red.

Since CHX and cold induction of *CBF2* is dependent on CAMTAs, we investigated whether these factors activate the cold transcriptome directly. Using epitope-tagged proteins complementing the *camta123* mutant (Supp. Fig. 3A,B), we performed ChIP-seq on CAMTA1, 2 and 3 in response to either cold or CHX treatment. Consistent with a direct role in activating early cold-responsive genes, the binding of these transcription factors is rapidly induced by both cold and CHX (Fig. 3C,D) and many of the genes in Cluster 1 are bound (Fig. 3C,F,G). CAMTAs also bind to the promoters of many genes in Clusters 3, 4 and 6, suggesting that they may also contribute to the regulation of late cold-responsive genes, which are strongly bound by CBF2 (Supp. Fig. 3C,D,E). We find 1141 genes where the binding of CAMTA1, 2 or 3 at their promoters increases during both cold and CHX treatments (Fig. 3E).

## Translational inhibition correlates with *CBF* gene induction

Since CHX has complex effects on the cell, we investigated whether its induction of the cold response is a specific consequence of its inhibition of ribosomes. Sampling a variety of translation inhibitors, we found a positive correlation between the degree of *CBF2* gene induction and the extent of translational inhibition (Fig. 4A). Inhibitors targeting different components of the translational machinery (Supp. Table 1) induce similar effects on the cold transcriptome, whereas terminating peptide elongation using puromycin has little effect (Fig. 4B).

**Figure 4:**
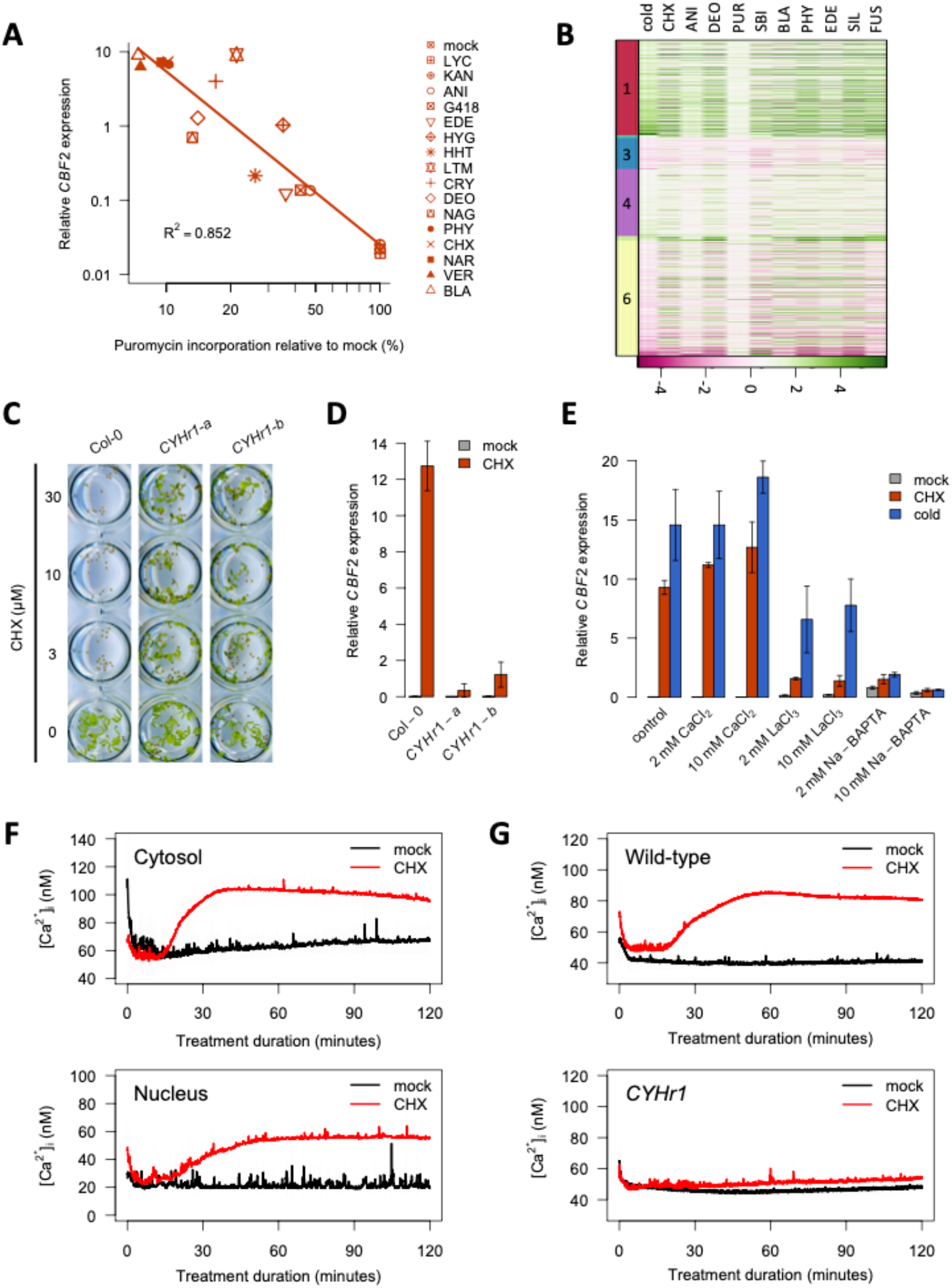
Ribosome-mediated cold gene activation involves calcium signalling. **A:** Translational activity correlates with *CBF2* gene expression. Seedlings were treated with inhibitors (30 μM, 22°C) or mock treatment (0.1% DMSO, 22°C) for 1 or 2 hours for translation assays and expression analyses, respectively. **B:** RNA-seq of seedlings after 2 hours of inhibitor, mock or cold (0.1% DMSO, 4°C) treatments. BLA: blasticidin S, ANI: anisomycin, LTM: lactimidomycin, HYG: hygromycin B, VER: verrucarin A, NAR: narciclasine, PHY: phyllanthoside, NAG: nagilactone C, DEO: deoxynivalenol, CRY: cryptopleurine, LYC: lycorine, HHT: homoharringtonine, EDE: edeine A1, KAN: kanamycin, SBI: SBI-0640756, SIL: silvestrol, FUS: fusidic acid. PUR, EDE and FUS treatments in (B) were at 150 μM as they do not induce *CBF2* expression at 30 μM. Colours indicate log2 fold-changes in expression relative to mock controls, for clusters from Fig. 2. **C:** Growth assay of wild-type and *CYHr1* seedlings in the presence of CHX or mock treatment. **D:** *CBF2* induction by CHX is abolished in *CYHr1* seedlings. Wild-type and *CYHr1* seedlings were treated with CHX or mock treatment for two hours. **E:** Calcium signalling inhibitors block *CBF2* induction by cold and CHX. CaCl2: calcium chloride, LaCl3: lanthanum chloride, Na-BAPTA: sodium BAPTA, control: media. Error bars indicate standard deviation for 3 replicates with 10-15 seedlings each (D, E). **F:** CHX triggers an increase in cytosolic and nuclear free calcium levels. **G:** The CHX-induced increase in intracellular calcium is abolished in *CYHr1* seedlings. Intracellular free calcium levels were quantified luminometrically in seedlings expressing apoaequorin in a localised (F) or ubiquitous (G) manner, during 2-hour CHX or mock treatments. Shading indicates standard deviation for at least three biological replicates, each comprising a cuvette with 3 seedlings.

To exclude the possibility that these chemicals activate the early cold response indirectly, we generated plants expressing the ribosomal protein point-mutant RPL36a^P56Q^ (*CYHr1,* for *cycloheximide resistance 1),* which in budding yeast does not interact with CHX (Kawai et al., 1992). *CYHr1* seedlings can grow in the presence of CHX (Fig. 4C), demonstrating conserved resistance conferred by the *RPL36a^P56Q^* mutation in plants. As expected, *CYHr1* plants are able to sustain protein synthesis when treated with CHX (Supp. Fig. 4), though some of ribosome inhibition still occurs as a result of two native *RPL36a* genes encoding CHX-binding proteins. *CYHr1* plants lose the ability to strongly induce *CBF2* gene expression in response to CHX, indicating that it is indeed the inhibition of ribosomal activity that transmits the cold signal (Fig. 4D).

## The cold signal is transmitted by a rise in cytosolic and nuclear calcium

Our results indicate that the rate of translation provides a direct readout of ambient temperature, and we are able to mimic cold-induced reductions in ribosomal activity by the use of chemicals, enabling the direct activation of early cold-responsive genes. This raises the question of how this signal is propagated in the cell. In maize, interfering with Ca^2+^ signalling inhibits the induction of cold-responsive genes by CHX (Berberich and Kusano, 1997). We were able to recapitulate these findings in *A. thaliana* using the Ca^2+^ signalling inhibitors lanthanum, a competitive inhibitor of Ca^2+^-permeable channels, and BAPTA, a Ca^2+^ chelator. These chemicals reduce the activation of *CBF2* expression in response to cold and CHX treatments (Fig. 4E). Similar reductions in cold-inducibility have been observed for other genes using Ca^2+^ signalling inhibitors in plants (Monroy et al., 1993; Knight et al., 1996; Polisensky and Braam, 1996; Tähtiharju et al., 1997).

Targeting the Ca^2+^ reporter aequorin to the nucleus or cytosol (Supp. Fig. 5), we found that CHX causes a rise in cytosolic and nuclear free Ca^2+^ (Fig. 4F). We obtained similar results using the Ca^2+^ reporter Cameleon localised in the cytosol (Supp. Fig. 6A, Supp. Movie). Since the induction of early cold-responsive genes by CHX is attenuated in *CYHr1* plants, we investigated changes in Ca^2+^ levels during CHX treatment in this line. No significant increase in intracellular free Ca^2+^ was detected in these plants (Fig. 4G). These results indicate that Ca^2+^ is a signal for activation of the early cold response during translation inhibition. CAMTA transcription factors are regulated by Ca^2+^-calmodulins (Bouché et al., 2005) and therefore provide a direct mechanism by which elevated nuclear free Ca^2+^ levels triggered by translation inhibition can activate early cold-responsive genes such as the *CBFs.* The 70S ribosome inhibitor kanamycin triggers a distinct cytosolic Ca^2+^ signature, but it does not induce *CBF* expression (Supp. Fig. 6B,C). This demonstrates that inhibiting cytosolic ribosomes generates a specific Ca^2+^ signal required for the activation of the early cold response.

Ribosome translation rates are also proportional to temperature in *E. coli* (Farewell and Neidhardt, 1998), and inhibiting translation in human cells also inactivates the heat shock response (Santagata et al., 2013), suggesting that ribosomes may have a broad role in providing thermosensory information. Since ribosomes must be active within the ambient temperature range of every organism, they are well suited to sense deviations from optimal temperature.

## Author contributions

DGS conceived the experiments, performed most of the experiments and wrote the first draft of the manuscript; KEJ performed the ChIP-seq experiments; FG performed the bioinformatic analysis in collaboration with DGS; FGD performed the Cameleon experiments; AC, AARW and PAW conceived the experiments and assisted in writing the manuscript.

## Plant materials and growth conditions

The *camta1, camta2, camta3, camta23* and *camta123* mutants were provided by Michael Thomashow (Kim *et al.,* 2013) and comprise the following T-DNA insertions: SALK_008187, SALK_007027 and SALK_001152. The *camta5, camta35, camta12346, camta12456*and *camta123456* mutants were provided by Kazuko Yamaguchi-Shinozaki (Kidokoro *et al.,* 2017) and include the following T-DNA insertions: SALK_108806, SALK_139868, SALK_001152, SALK_087870, SALK_134491 and SALK_078900. The *prr579* mutant *(prr5-11 prr7-11 prr9-10)* was provided by Norihito Nakamichi (Nakamichi *et al.,* 2005) and consists of T-DNA insertions SALK_064538, SALK_030430 and SALK_007551. The Col-0 *pCaMV35S::APOAEQUORIN* line was provided by Alex Webb (Xu *et al.,* 2007), and the Col-0 *pUBQ10-NES::YC3.6* line was provided by Melanie Krebs (Krebs *et al.,* 2012).

Seeds were surface-sterilised using ethanol or vapour-phase sterilisation (Clough & Bent, 1998) and stratified in respective growth media in the dark at 4°C for 72 hours. Growth cabinets were maintained at 65% relative humidity and 170 μmol/m^2^/s light, unless specified otherwise, and as specified per experiment seedlings were grown at 20°C with a 12-hour photoperiod or at 22°C with either continuous light, a 16-hour long-day photoperiod or an 8-hour short-day photoperiod.

For ChIP-seq, *A. thaliana* seedlings were grown for 8 days on 1-mm-pore nylon mesh rafts placed on half-strength MS medium (pH 5.7) with 0.8% w/v agar (P1001, Duchefa) (‘MS agar’ hereafter) (22°C long days).

For Cameleon experiments, seedlings were grown for 6-7 days on vertical plates of MS agar supplemented with 0.1% w/v sucrose and 0.05% w/v MES (22°C long days, with Cool White Neon lamps at 100 μmol/m^2^/s).

For gene expression analyses, SUnSET assays, seedlings were grown for 7-9 days in half-strength Murashige-Skoog liquid medium (½×MS; 0.22% w/v Murashige-Skoog mix including vitamins; M0222, Duchefa) containing 0.05% w/v MES (2-[N-morpholino]ethanesulfonic acid; 69892, Sigma), adjusted to pH 5.7 and supplemented with 0.1% w/v glucose after autoclaving (‘½MMG medium’ hereafter) (22°C long days, with the following exceptions: 22°C short days for Fig. 1C, and 22°C continuous light for Figs. 2B, 3A, 3B, 4E). The seedlings were cultured in 12- or 6-well plates (657-160 and 665-180, Greiner Bio-One), with 1 mL or 2 mL of liquid medium per well, respectively, and sealed with Micropore tape. For aequorin-based calcium measurements, seedlings were grown in ½MMG for 8 to 12 days at 20°C with a 12-hour photoperiod.

For transformations and seed harvesting, *A. thaliana* plants were grown on Levington F2 soil (22°C long days). Transgenic *A. thaliana* T1 and T2 seeds were stratified and grown for selection on MS agar supplemented with 50 to 100 μg/mL kanamycin.

## Temperature and chemical treatments

Inhibitors were prepared as indicated in *Table 1.* Edeine A1 was provided by Ian Brierley (University of Cambridge) and phyllanthoside, cryptopleurine, narciclasine, nagilactone C and silvestrol were provided by the Developmental Therapeutics Program of the National Cancer Institute (NCI), National Institute of Health (NIH). Chemical treatments were performed by replacing the liquid medium with fresh ½MMG medium containing chemicals diluted to the specified concentrations. Cold shock treatments were performed by transferring plates to 4°C pre-cooled cabinets, at 170 μmol/m^2^/s light and 65% relative humidity.

**Table 1:** Inhibitors used in this study

| Chemical | Target | Mechanism of inhibition | Supplier | Catalogue number | Stock solvent |
| --- | --- | --- | --- | --- | --- |
| Anisomycin | A-site (60S subunit) | Peptide bond formation | Sigma | A9789 | DMSO |
| BAPTA | - | Ca<sup>2+</sup> chelation | Abcam | ab120449 | - |
| Blasticidin S | P-site (60S/50S subunits) | Peptide bond formation, termination | Sigma | 15205 | DMSO |
| Cryptopleurine | mRNA tunnel (40S subunit) | Translocation | NIH/NCI | NSC 19912 | DMSO |
| Cycloheximide | E-site (60S subunit) | Translocation | Sigma | 46401 | DMSO |
| Deoxynivalenol | A-site (60S subunit) | Peptide bond formation | Sigma | D0156 | DMSO |
| Edeine A1 | mRNA tunnel (40S/30S subunits) | Initiation | Ian Brierley (Cambridge) | - | water |
| Fusidic acid | Elongation factors | GTPase | Sigma | F0881 | water |
| G418 | Decoding centre (40S/30S subunits) | Translocation, aminoacyl-tRNA selection | Sigma | A1720 | water |
| Homo-harringtonine | A-site (60S subunit) | Peptide bond formation | Sigma | SML1091 | DMSO |
| Hygromycin B | Decoding centre (40S/30S subunits) | Translocation, aminoacyl-tRNA selection | Sigma | H9773 | DMSO |
| Kanamycin | A-site (30S subunit) | Translocation, aminoacyl-tRNA selection | Fisher | BP906-5 | water |
| Lactimidomycin | E-site (60S subunit) | Translocation | Merck Millipore | 506291 | DMSO |
| Lanthanum | - | Ca<sup>2+</sup> channel blocker | Sigma | 262072 | - |
| Lycorine | A-site (60S subunit) | Peptide bond formation | Sigma | L5139 | DMSO |
| Nagilactone C | A-site (60S subunit) | Peptide bond formation | NIH/NCI | NSC 211500 | DMSO |
| Narciclasine | A-site (60S subunit) | Peptide bond formation | NIH/NCI | NSC 266535 | DMSO |
| Phyllanthoside | E-site (60S subunit) | Translocation | NIH/NCI | NSC 328426 | DMSO |
| Puromycin | A-site (60S/50S subunits) | Triggers termination | Sigma | P8833 | DMSO |
| SBI-0640756 | eIF4G | Initiation | Sigma | SML1645 | DMSO |
| Silvestrol | eIF4A | Initiation | NIH/NCI | NSC 783538 | DMSO |
| Verrucarin A | A-site (60S subunit) | Peptide bond formation | Sigma | V4877 | DMSO |

For ChIP-seq samples, plants on MS agar were incubated at 4°C for four hours (CBF2 ChIP-seq) or submerged with 30 μM CHX or 0.1% v/v DMSO (mock control) in deionised water and maintained at 22°C or 4°C for one or two hours (CAMTA ChIP-seq).

For SUnSET assays to measure *in vivo* translation (Schmidt *et al.,* 2009), following temperature or chemical treatments, the media was supplemented with either 100 or 150 μM puromycin (in ½MMG; temperature-adjusted) and seedlings were harvested after 20 or 30 minutes of incubation.

For growth assays of Col-0 and *CYHr1* seedlings in the presence of CHX, the ½MMG media was removed after two days of growth and replaced with fresh media containing CHX or DMSO to final concentrations of 3, 10 or 30 μM or 0.1% v/v, respectively, and seedlings were imaged after seven days of growth (22°C long days).

## Generation of transgenic lines

All vectors were constructed using Ligation-Independent Cloning, as described by Li & Evans (1997), using a 30-second digestion with ExoIII, and were transformed into *A. thaliana* using the floral dip method (Clough & Bent, 1998). Primers used for cloning are given in *Table 2.*

**Table 2:**
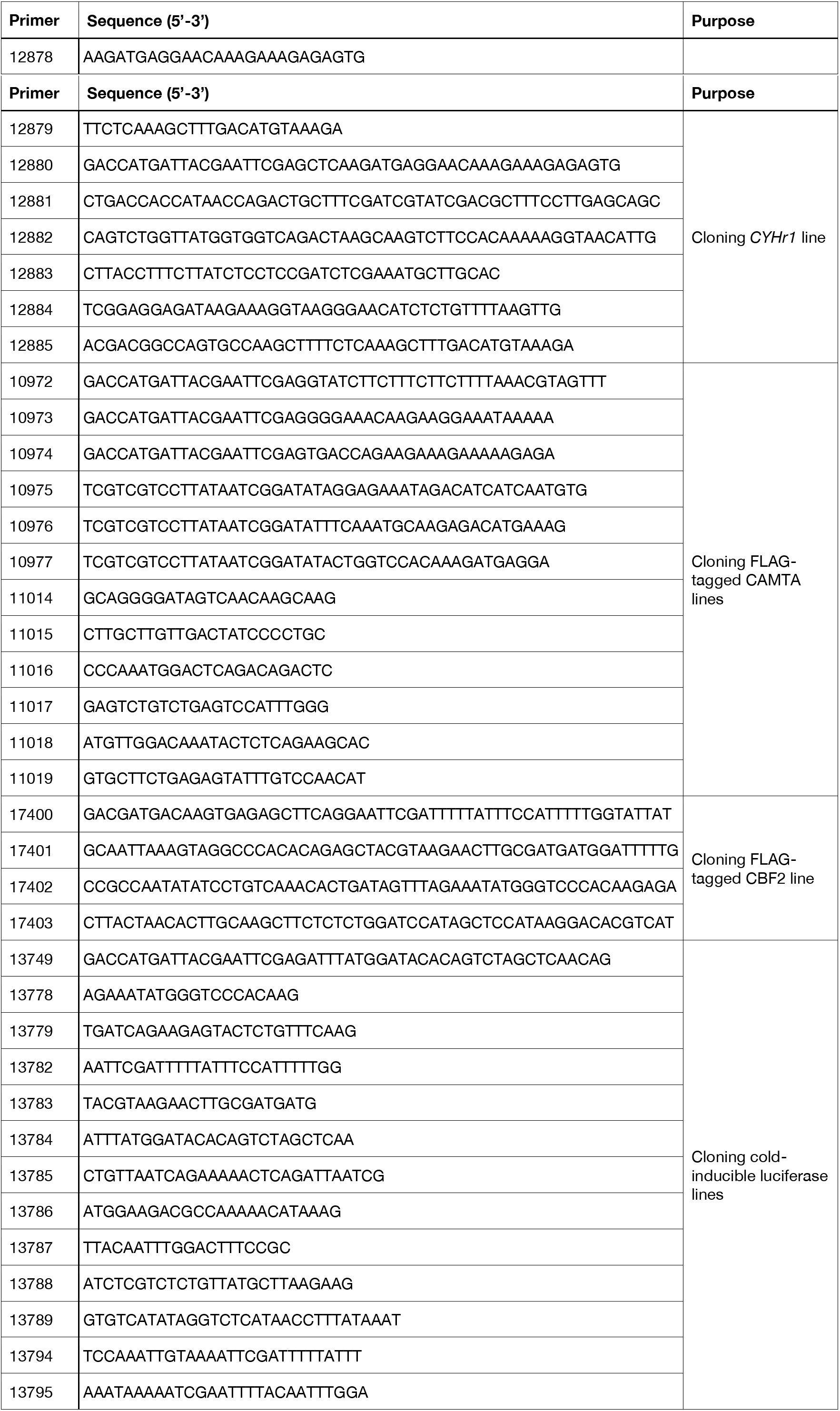
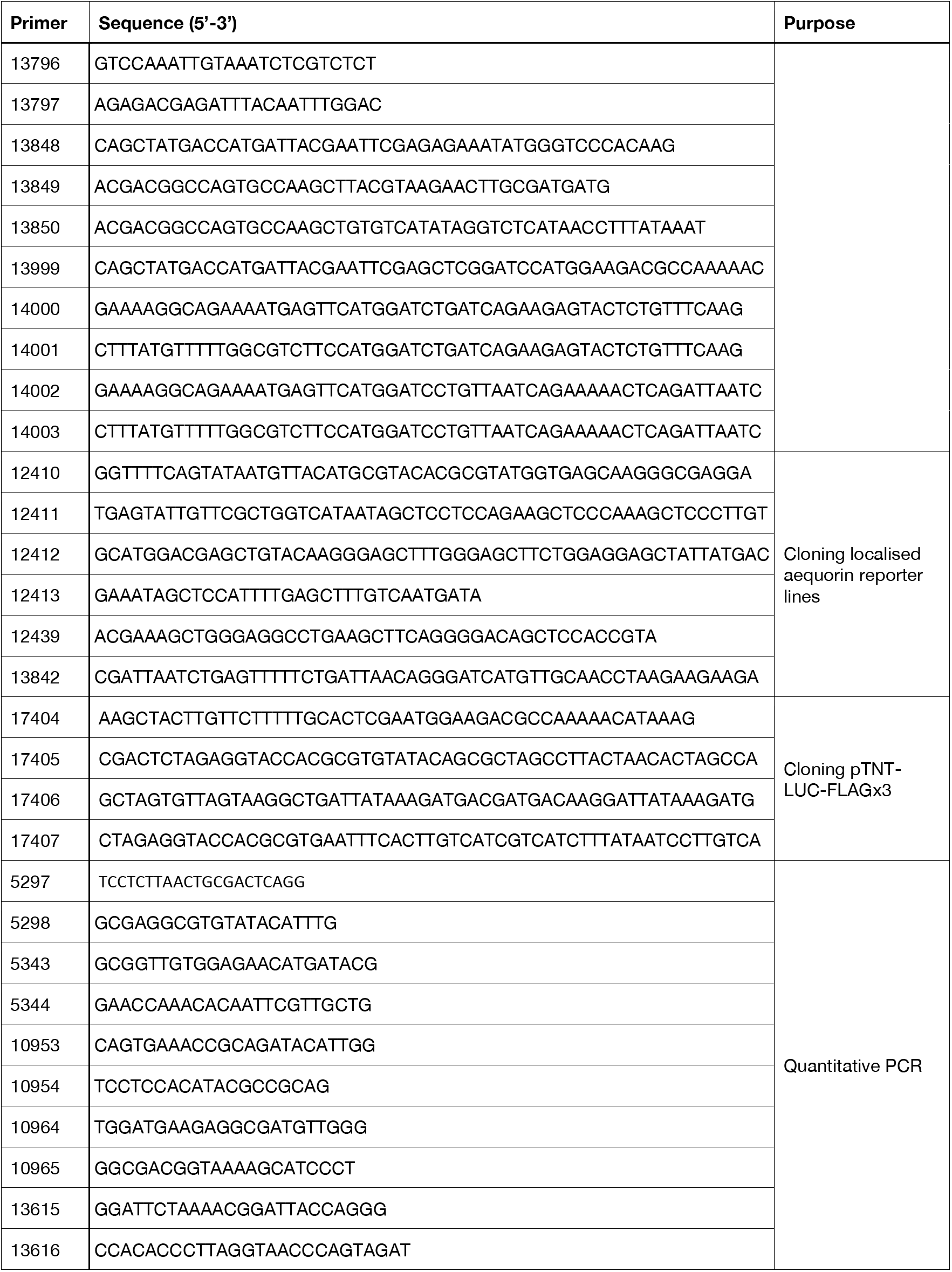
Primers used in this study

CHX-resistant *CYHr1* line: the *RPL36aA* gene was amplified from Col-0 genomic DNA with primers 12878+12879. Mutated fragments were generated with primers 12880+12881, 12882+12883 and 12884+12885 and were joined by overlap PCR to produce the gene *RPL36aAΔP56Q.* The binary vector *PW1211* (Philip A. Wigge, unpublished) was linearised with Eco53kI and HindIII, and both *RPL36aA^P56Q^* and *PW1211* were recombined to produce *pCYHr1*

*(pRPL36aA::RPL36aA^P56Q^::tRPL36aA),* which was subsequently transformed into Col-0 and Col-0 *pCaMV35S::APOAEQUORIN.* The mutated *RPL36aA* gene was expressed under its native promoter and terminator sequences. Two independent transgenic lines were used for experiments, named *CYHr1-a* and *CYHr1-b.*

CAMTA tagged lines for ChIP-seq: the *CAMTA1, CAMTA2* and *CAMTA3* genes were amplified in two parts from Col-0 genomic DNA with primers 10972+11015 and 11014+10975, 10973+11017 and 11016+10976, and 10974+11019 and 11018+10977, respectively. *PW1211* was linearised with Eco53kI and EcoRV and recombined with both inserts to produce the plasmids *pCAMTA1::CAMTA1::FLAG×3, pCAMTA2::CAMTA2::FLAG×3* and *pCAMTA3::CAMTA3::FLAG×3,* which were subsequently transformed into the *camta123* triple mutant (Kim *et al.,* 2013). All FLAG-tagged CAMTAs were expressed under their native promoter and terminator sequences.

CBF2 tagged line for ChIP-seq: the *CBF2* terminator was amplified from Col-0 genomic DNA with primers 17400+17401 and recombined with the vector *pUBQ10-CFLAG* (David Guillaume-Schoepfer & Philip A. Wigge, unpublished), linearised with StuI and AfeI, to produce the plasmid *p3FLAG-tCBF2*. The *CBF2* gene was amplified with primers 17402+17403 and recombined with *p3FLAG-tCBF2,* linearised with PmeI and BamHI, to generate the plasmid *pCBF2::CBF2::FLAGx3::tCBF2,* which was subsequently transformed into Col-0. FLAG-tagged CBF2 was expressed under its native promoter and terminator sequences.

Cold-inducible luciferase reporter lines: the *LUC* coding sequence was amplified from *pBGWL7* (Karimi *et al.,* 2005) with primers 13786+13787, and the *CBF2* and *UBQ10* promoters and terminators were amplified from Col-0 genomic DNA with primers 13778+13779, 13784+13785, 13782+13783 and 13788+13789, respectively. Fragments were re-amplified with primers 13999+13794 or 13999+13796 (*LUC*), 13795+13849 *(tCBF2),* 13797+13850 *(tUBQ10),* 13848+14000 or 13848+14001 *(pCBF2)* and 13749+14002 or 13749+14003 *(pUBQ10),* to produce overlapping ends, and the *LUC* coding sequence was joined to terminator regions by overlap PCR. *PW1211* was linearised with Eco53kI and HindIII and recombined with the overlapped amplicons. The resultant vectors were linearised with Eco53kI and BamHI and recombined with promoter regions to produce the plasmids *pCBF2::LUC::tCBF2*, *pCBF2::LUC::tUBQ10, pUBQ10::LUC::tUBQ10* and *pUBQ10::LUC::tCBF2,* which were transformed into Col-0.

Localised aequorin reporter lines: the coding sequences of *VENUS* and *APOAEQUORIN* were amplified from *P2R-P3a_4glyVenusYFP-3AT* (Yrjö Helariutta, unpublished) and *pNEWAEQ* (Alex Webb, unpublished) vectors using primers 12410+12411 and 12412+12413, respectively, and were joined by overlap PCR. The *VENUS::APOAEQUORIN* amplicon was re-amplified with primers 13842+12439 and 13843+12439 to introduce the SV40 nuclear localisation sequence or the human PK1α nuclear export sequence, respectively (Mehlmer *et al.,* 2012). These fragments were recombined with the vector *pUBQ10-3AT* (David Guillaume-Schöpfer & Philip A. Wigge, unpublished), linearised with BamHI and HindIII, to produce the plasmids *pUBQ10::NLS_SV40_::VENUS::APOAEQUORIN* and *pUBQ10::NES_PK1α_::VENUS::APOAEQUORIN,* which were subsequently transformed into Col-0.

Plasmid for *in vitro* transcription and translation: the coding sequence of *LUC* was amplified from *pBGWL7* with primers 17404+17405 and recombined with the vector *pTNT* (L5610, Promega), linearised with XhoI and EcoRI. The resultant plasmid was linearised with AfeI and BstZ17I and recombined with primers 17406+17407, annealed to produce the FLAG tag, to generate the plasmid *pTNT-LUC-FLAG×3.*

## Gene expression analyses

For expression analyses by quantitative PCR, total RNA was isolated from *A. thaliana* seedlings using phenol-chloroform extraction, as described by Box *et al.* (2011). Genomic DNA was eliminated by DNase I treatment using the TURBO DNA-free kit (AM1907, Invitrogen) and purified RNA was reverse-transcribed using the Transcriptor First-Strand cDNA Synthesis Kit (04379012001, Roche), according to manufacturer instructions. Quantitative real-time PCR was carried out using a LightCycler 480 (Roche) with DNA SYBR Green I Master mix (04707516001, Roche). The following primers were used for quantitative PCR, with sequences listed in *Table 2:* 10964+10965 *(CBF2/AT4G25470),* 5343+5344 *(PP2A/AT1G13320),* 5297+5298 *(UBC21/AT5G25760),* 10953+10954 *(COR15a/AT2G42540)* and 13615+13616 (*LUC*). The expression of *CBF2, COR15a* and *LUC* was normalised to the expression of both *PP2A* and *UBC21.* Because of the similarity in sequence between *CBF1, CBF2* and *CBF3* genes, the *‘CBF2’* primers used give an indication of total *CBF* expression. Crossing point (Cp) values were calculated using the ‘Second Derivative Maximum Method’ in the LightCycler software, and relative gene expression was subsequently calculated using the following formula:

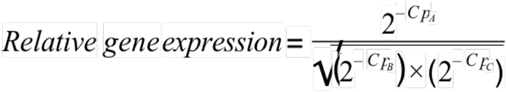

where *A* is the gene of interest and *B* and *C are* control genes used for normalisation.

For transcriptomic analyses by RNA-seq, total RNA was extracted and DNase-treated using MagMAX-96 Total RNA Isolation Kit (AM1830, Thermo Fisher Scientific). RNA was quantified using the Qubit fluorometer (Thermo Fisher Scientific) and RNA profiles were analysed using a TapeStation 2200 (Agilent) with RNA ScreenTapes (5067-5576, Agilent). Libraries for sequencing were prepared from 500 ng RNA using the QuantSeq 3’ mRNA-Seq Library Prep Kit (Lexogen), according to manufacturer instructions. DNA libraries were quantified using a Qubit fluorometer, and library profiles were analysed using a TapeStation 2200 with High Sensitivity D1000 ScreenTapes (5067-5584, Agilent). Libraries were sequenced on a NextSeq-500 (Illumina; paired-end 75bp reads), according to Illumina guidelines.

## Chromatin immunoprecipitation

The seedling were grown for 10 days and treated as indicated. 3 g plant material for each treatment was fixed under vacuum for 20 min in 1xPBS (10 mM PO_4_^3-^, 137 mM NaCl, and 2.7 mM KCl) containing 1% Formaldehyde (F8775 SIGMA). The reaction was quenched by adding glycine to a final concentration of 62 mM. Chromatin immunoprecipitation (ChIP) was performed as described (Jaeger et al), with the exception that 100 μl of ANTI-FLAG® M2 Affinity Gel (A2220 Sigma) were used for Immunoprecipitation seedlings. Sequencing libraries were prepared using TruSeq ChIP Sample Preparation Kit (Illumina IP-202-1024) and prepared according to manufacturer instructions. DNA libraries were quantified using a Qubit fluorometer, and library profiles were analysed using a TapeStation 2200 with High Sensitivity D1000 ScreenTapes (5067-5584, Agilent). Libraries were sequenced on a NextSeq-500 (Illumina; single end, 75bp reads), according to Illumina guidelines.

## Bioinformatic methods

RNA-seq samples were analysed with a commercial pipeline (QuantSeq FWD; Bluebee).

From the processed count files, CPM values (counts per million reads mapped) were calculated for each annotated gene and used as the abundance measure.

For ChIP-seq reads, adaptor contamination and low-quality trailing sequences were removed using Trimmomatic (Bolger *et al.,* 2014). Trimmed reads were mapped to the TAIR10 transcriptome using Bowtie2 (Langmead & Salzberg, 2012). Any read that mapped to more than one genomic location was discarded and optical duplicates were removed using Picard (http://github.com/broadinstitute/picard). Genomic binding profiles were quantified in RPKM (reads per kilobase per million mapped reads) using a bin-size of 10 bp. Peaks were identified with Model-based Analysis of ChIP-seq (MACS2) (Zhang *et al.,* 2008) with argument “--keep-dup 1 -p 0.1”, and peaks were filtered for fold-change > 4 (Fig. 3) or > 6 (Supp. Fig. 3E). Any gene containing a peak summit within 3 kb of its start codon was classified as a bound target. For ChIP-seq pile-ups, RPKM profiles were extracted for each peak around the MACS2-reported summit position and per-position averages and standard deviations were calculated across target peaks. The ChIP-seq data was visualised with the Integrated Genome Viewer (IGV), and for IGV snapshots the same y-axis scales were used for all tracks (Fig. 3C and Supp. Fig. 3D).

Gene lists for Venn diagrams, top 1000 high-confidence (based on FC) ChIP genes Raw and processed RNA-seq and ChIP-seq data are available online (NCBI Geo Ominibus accession GSEXXXXXX and GSEXXXXXX). Code for reproducing the analysis is available at https://www.github.com/shouldsee/camta-figures.

Analyses of promoter motif enrichment were carried out using Homer2 (Heinz *et al.,* 2010) with promoter sequences retrieved from TAIR (TAIR10 Loci Upstream Seq −1000bp; https://www.arabidopsis.org/tools/bulk/sequences/).

## In vitro transcription and translation

*pTNT-LUC-FLAG×3,* described above, was linearised with BamHI and uncapped transcripts were synthesised *in vitro* for 6 hours at 37°C using bacteriophage T7 RNA polymerase (EP0112, Thermo Fisher Scientific) in a 50 μL reaction supplemented with 20 U SUPERase-In RNase inhibitor (AM2694, Thermo Fisher Scientific), 4 mM NTPs and 5 mM DTT. DNA was removed in a 30-minute digestion with DNAse I at 37°C and RNA was subsequently purified using 1:1 phenol-chloroform (pH 4.3) and ethanol-precipitated.

*In vitro* translation was performed for one hour at the temperatures indicated (0°C to 35°C), using wheat germ extracts (L4380, Promega) in 40 μL reactions supplemented with all amino acids, 70 mM potassium acetate, 0.4 ug denatured RNA, 16 U SUPERase-In RNase inhibitor (AM2694, Thermo Fisher Scientific) and 1*×* complete EDTA-free protease inhibitors (11873580001, Roche). Reactions were stopped by adding CHX and EDTA to final concentrations of 100 μM and 300 μM, respectively, and placing the tubes on ice. To measure the activity of recombinant luciferase, sodium-D-luciferin and ATP were added to the reactions to final concentrations of 100 μM and 50 μM, respectively, and luminescence was quantified at room temperature using a TriStar LB-942 plate reader (Berthold). To measure the protein yield, reactions were mixed with SDS loading buffer (60 mM Tris-HCl pH 6.8, 10% v/v glycerine, 2% w/v SDS, 100 mM DTT, 0.015% w/v bromophenol blue; final concentrations) and FLAG-labelled proteins were analysed as described below.

## Western blotting

Frozen seedlings were pulverised in 2 mL tubes containing a single tungsten carbide bead (69997, Qiagen) using the Tissue Lyser II (Qiagen). Proteins were extracted using the urea-SDS method outlined by Clontech (2009), with 150 μL cracking buffer per 10 to 20 pulverised seedlings. Protein extracts were separated by SDS-polyacrylamide gel electrophoresis, transferred onto PVDF membranes and probed with horseradish peroxidase (HRP)-conjugated anti-FLAG (A8592, Sigma; 1:3000), anti-puromycin (MABE343, Sigma; 1:1000), anti-actin (A0480, Sigma; 1:3000) or anti-RPS14 (AS12-2111, Agrisera; 1:3000) primary antibodies and DyLight 800-conjugated anti-mouse IgG (AS12-2426, Agrisera) or DyLight 650-conjugated anti-rabbit IgG (AS12-2327, Agrisera) secondary antibodies. Washes were carried out in 1 *×* TBST buffer (20 mM Tris-HCl, 150 mM NaCl, 0.1% v/v Tween-20) and blocking and incubations were carried out with 5% w/v BSA or skimmed milk in 1 *×* TBST buffer. Chemiluminescence was quantified using Pierce ECL Western Blotting Substrate (32106, Thermo Scientific) over a period of five minutes, and 600 nm, 700 nm or 800 nm fluorescence emission detection was measured for 30 seconds, using a dual-mode camera (Odyssey Fc, Li-Cor) coupled with an imaging suite (Image Studio v.2.1.10, Li-Cor). The intensity of protein bands was measured using ImageJ (NIH).

For SUnSET assays (Schmidt *et al.,* 2009), the level of *in vivo* translation was determined by measuring the amount of puromycin-labelled proteins, normalised to actin levels, and is given as a percentage of maximal puromycin incorporation. For time-course experiments, puromycin incorporation in CHX- and cold-treated seedlings were additionally normalised to that in mock controls for each time point.

## Fluorescence microscopy

Confocal images were obtained using a Leica SP8 laser-scanning confocal microscope with 20*×*/0.75 or 63*×*/1.2 objectives (Leica Microsystems, Wetzlar, Germany). The fluorescent protein Venus (modified yellow fluorescent protein, YFP) was excited using a white light laser at 514 nm, with detection restricted to 520-550 nm.

Samples for immunofluorescence microscopy were prepared as described by Pasternak *et al.* (2015), with the following modifications. Seven-days-old seedlings were fixed for two hours in 2% formaldehyde solution and, following tissue clearing with methanol, were digested for 8 minutes at 37°C in digestion solution (4 ng/μL cellulase R-10 [C8001, Duchefa], 4 ng/μL macerozyme R-10 [M8002, Duchefa], 6 ng/μL pectinase [17389, Sigma-Aldrich], 2 ng/μL pectolyase Y-23 [P8004, Duchefa] in 1*×* PBST buffer [137 mM NaCl, 2.7 mM KCl, 8 mM Na2HPO4, 1.5 mM KH2PO4, 0.01% v/v Tween-20, pH 4.7]). Following membrane permeabilisation, seedlings were incubated with gentle mixing firstly in blocking solution (5% BSA in 1*×* PBST buffer) for 2 hours at room temperature and then in 1:500 antibody solutions (anti-puromycin [MABE343, Sigma] primary antibody, and DyLight 650-conjugated anti-mouse IgG [AS12-2302, Agrisera] secondary antibody, both in blocking solution) overnight at 4°C, with three washes with 1 *×* PBST buffer after each antibody incubation. DyLight 650 was excited using a white laser at 633 nm, with detection restricted to 668-678 nm. Images were analysed with ImageJ (NIH).

For wide-field Ca^2+^ imaging analyses in *A. thaliana* root tip cells, an inverted fluorescence Nikon microscope (Ti-E) with a 20× N.A. 0.75 was used. Excitation light was produced by a fluorescent lamp (Prior Lumen 200 PRO, Prior Scientific) set to 20% with 440 nm (436/20 nm) excitation for the Cameleon (YC3.6) sensor. Images were collected with a Hamamatsu Dual CCD camera (ORCA-D2). The FRET CFP/YFP optical block A11400-03 (emission 1, 483/32 nm for CFP; emission 2, 542/27 nm for FRET) was used with a dichroic 510-nm mirror (Hamamatsu) for simultaneous CFP and cpVenus acquisitions. Camera binning was set to 2 × 2 with exposure times of 200 ms. Images where acquired every 5 s. Filters and dichroic mirrors were purchased from Chroma Technology. NIS-ElementsTM (Nikon) was used as a platform to control the microscope, illuminator and camera. Images were analysed using FIJI.

## Measurements of intracellular free calcium

For aequorin-based measurements, three seedlings grown in ½MMG medium for eight to twelve days were transferred to 500 μL freshly-prepared 2 μM coelenterazine solution (303-500, Nanolight Technology; in deionised water with 0.5% v/v methanol) in luminometer cuvettes (diameter 12 mm, height 51 mm; Sarstedt). Following reconstitution of aequorin in the dark at 20°C for at least 10 hours, the coelenterazine was replaced with 500 μL 30 μM CHX, 30 μM kanamycin or 0.1% v/v DMSO (mock control) in deionised water at 20°C. The cuvette was immediately placed in a photon-counting luminometer (9899A photomultiplier tube; cooled to −20°C with a FACT50 housing [Electron Tubes]) and photon counts were measured every second. To estimate total aequorin in the samples, 1 mL of discharge solution (1 M CaCl2, 10% v/v ethanol, final concentrations) was injected into the cuvette through a light-tight port in the luminometer using a 1 mL syringe and 75 mm needle. Measurements were continued until photon counts/sec had reached <10% of the discharge peak. Nuclear and cytosolic free calcium levels were measured using *pUBQ10::NLS_SV40_::VENUS::APOAEQUORIN* and *pUBQ10::NES_PK1α_::VENUS::APOAEQUORIN* (described above), respectively. Col-0 *pCaMV35S::APOAEQUORIN* and *CYHr1 pCaMV35S::APOAEQUORIN* were used for non-localised measurements of intracellular free calcium ions (cytosolic and nuclear). Calcium concentrations were estimated according to Fricker *et al.* (1999) using the following formula:

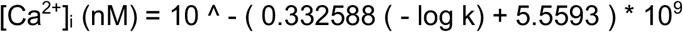

where k = photon count / total photon count over course of experiment.

For Cameleon-based measurements, seedlings grown on MS agar for seven days were gently transferred to dedicated chambers and overlaid with cotton wool soaked in imaging solution (5 mM KCl, 10 mM MES, 10 mM CaCl2, adjusted to pH 5.8 with Tris). The root was continuously perfused with imaging solution, whereas the shoot was not submerged. The treatment was carried out by adding CHX to the imaging solution at a final concentration of 30 μM at the indicated time. Fluorescence intensity was determined over a region of interest (ROI), corresponding to the root tip meristematic zone (Behera *et al.,* 2018). cpVenus and CFP emissions in the analysed ROI were used for the ratio (R) calculation (cpVenus/CFP), which was normalised to the initial ratio (R_0_) and plotted versus time (ΔR/R_0_). Background subtraction was performed independently for both channels before calculating the ratio.

---

**Supplementary Figure 1:**
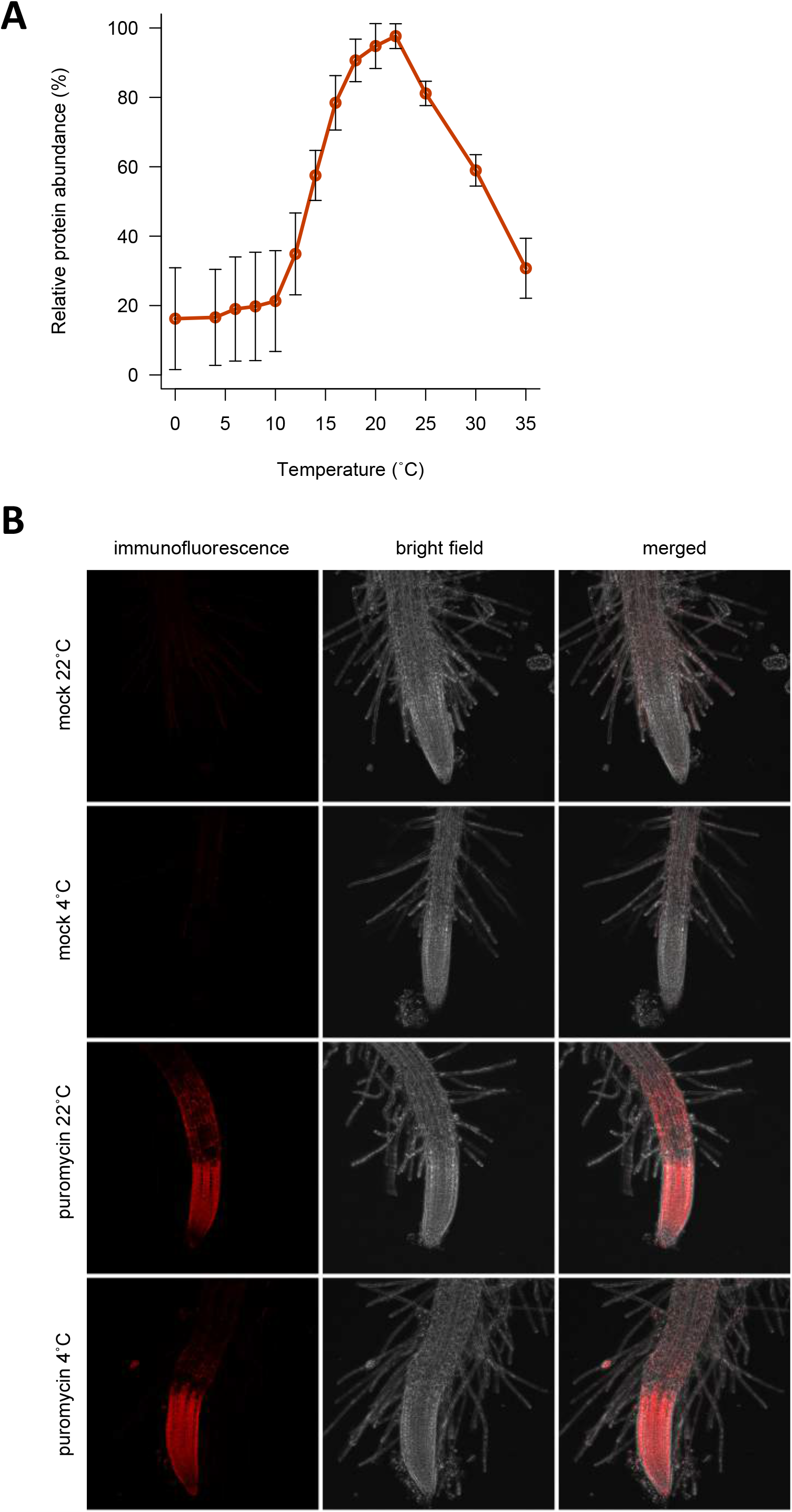
Ambient temperature has an inherent effect on translation. **A:** Low temperatures reduce protein synthesis *in vitro.* Protein yield of C-terminally FLAG-tagged luciferase synthesised *in vitro* using wheat germ extract for 1 hour at 0°C to 35°C. FLAG-labelled protein levels were normalised to RPS14 levels and are given as a percentage of the maximum yield. All translation reactions contained equal amounts of mRNA. Error bars indicate the standard deviation for 4 replicate reactions. **B:** Low temperatures do not prevent the cellular uptake of puromycin. *Arabidopsis thaliana* Col-0 seedlings were grown in liquid culture in long days at 22°C for 7 days and, following a 2-hour pre-incubation at either 4°C or 22°C, were treated with 150 μM puromycin or 0.1% DMSO (mock) for 30 minutes at these respective temperatures and subsequently fixed with formaldehyde. Intracellular puromycin was detected by immunofluorescence microscopy for 7 biological replicates with the same laser settings and representative images are shown.

**Supplementary Figure 2:**
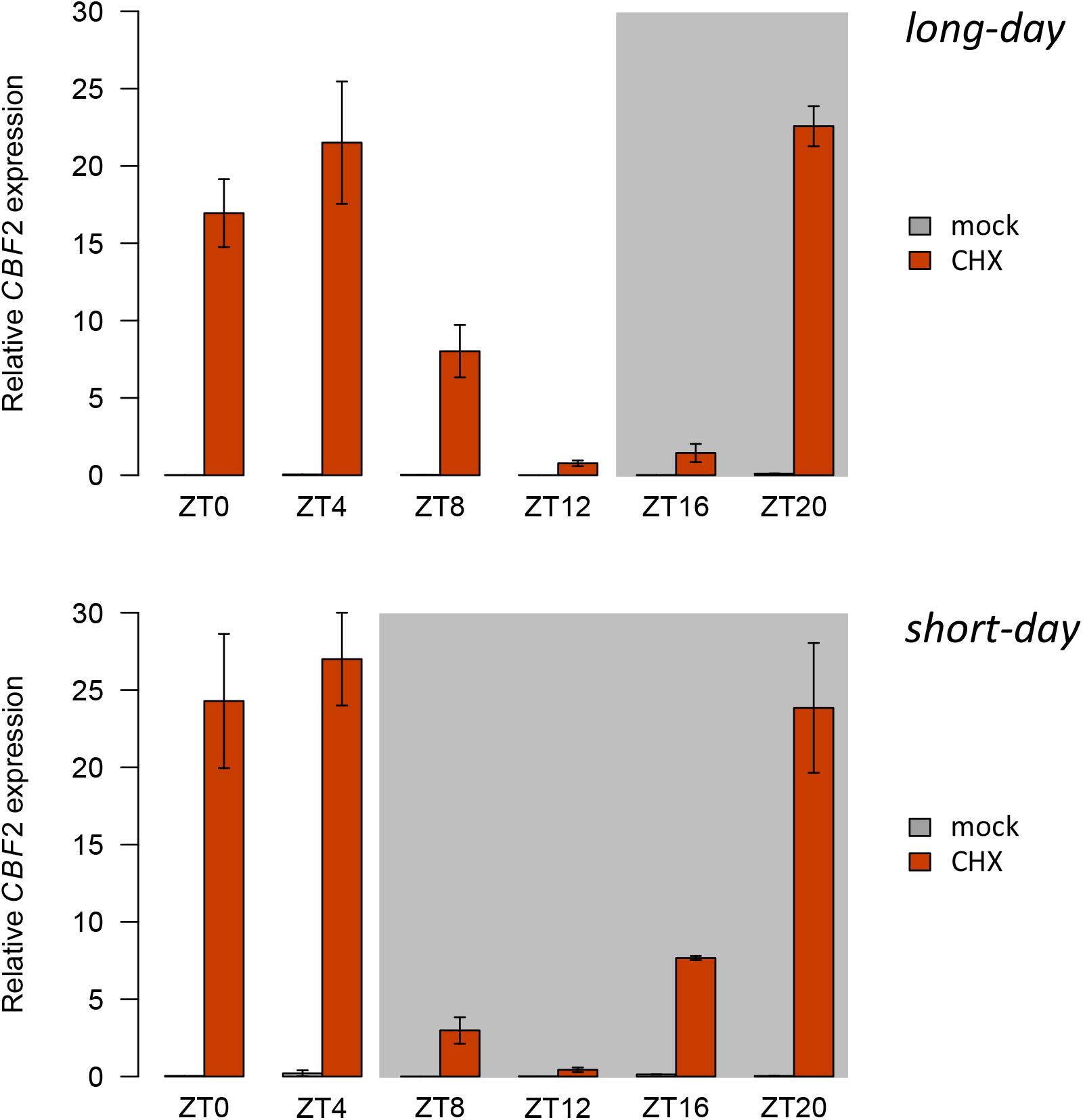
*CBF2* induction by CHX treatments varies across the 24-hour cycle. *A. thaliana* Col-0 seedlings were grown in liquid culture for 9 days at 22°C with a long-day (16-hour) or short-day (8-hour) photoperiod and harvested after 2-hour treatments at 22°C with 30 μM CHX or 0.1% DMSO (mock) at the indicated intervals. *CBF2* expression was determined by quantitative PCR and normalised to transcript levels of *PP2A* and *UBC21.* Error bars represent standard deviation for 3 biological replicates, with 10-15 seedlings per replicate. Grey boxes represent treatments performed in the dark.

**Supplementary Figure 3:**
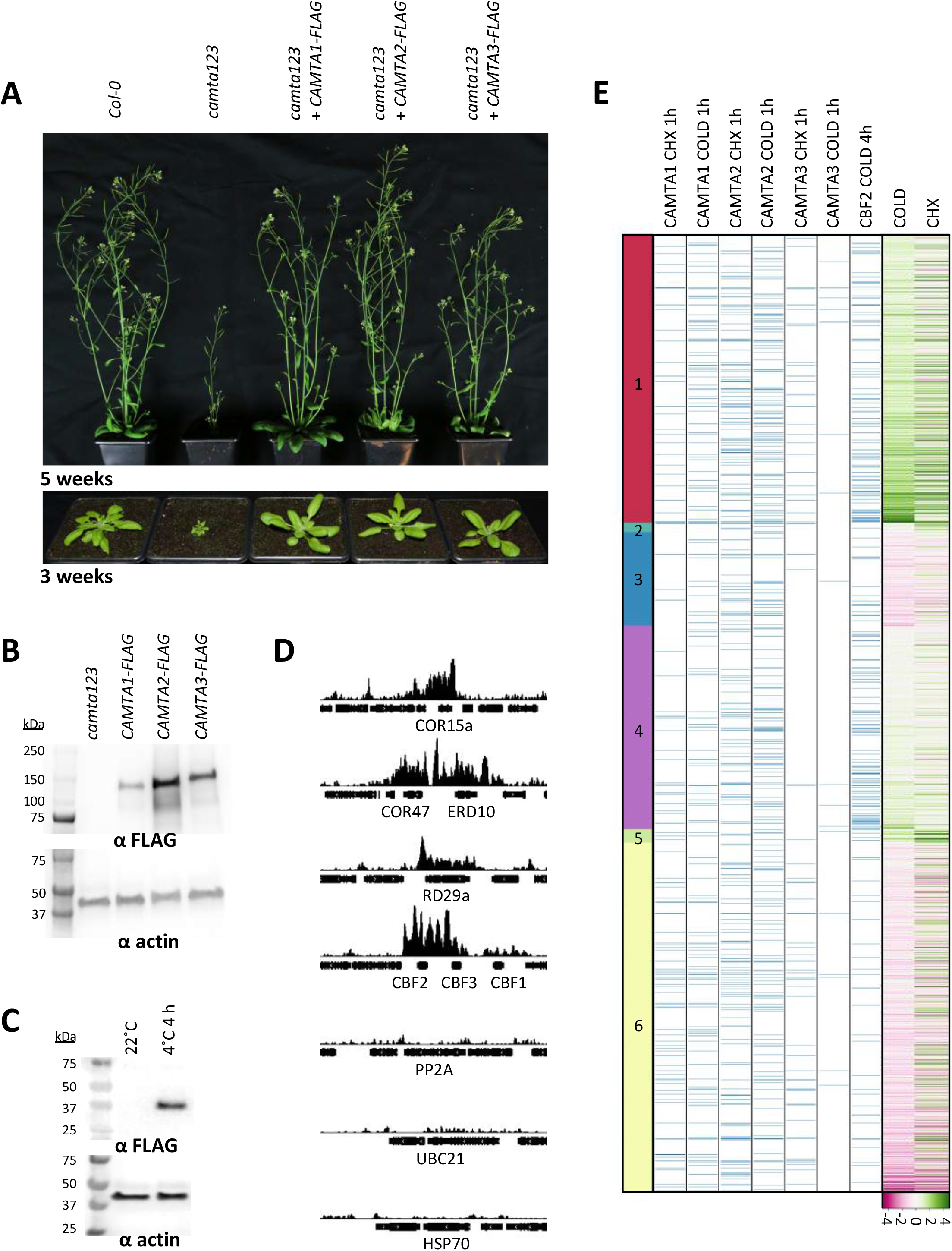
ChIP-seq of FLAG-tagged lines. **A:** Complementation of the *camta123* mutant with a C-terminal FLAG-tagged CAMTA1, 2 or 3 *(pCAMTA1::CAMTA1::FLAG×3, pCAMTA2::CAMTA2::FLAG×3,* or *pCAMTA3::CAMTA3::FLAG× 3).* Images of homozygous *A. thaliana* plants after 3 or 5 weeks of growth at 22°C in long days. **B:** Protein expression of FLAG-tagged CAMTA1, 2 and 3, and of actin as a loading control, in homozygous complemented camta123 *A. thaliana* seedlings grown for 7 days in liquid medium at 22°C in long days. **C:** Protein expression of FLAG-tagged CBF2, and of actin as a loading control, in homozygous Col-0 seedlings grown for 7 days in liquid medium at 22°C in long days and either transferred to 4°C for 4 hours or maintained at 22°C. **D:** Cold-associated binding of CBF2 at coldinducible genes *COR15a* (AT2G42540), *COR47* (AT1G20440), *ERD10* (AT1G20450), *RD29a* (AT5G52310), *CBF2* (AT4G25470) and *CBF3* (AT4G25480), 4 hours after transfer from 22°C to 4°C. No binding is detected at control genes *PP2A* (AT1G13320), *UBC21* (AT5G25760) and *HSP70* (AT3G12580), which are not induced at low temperatures. **E:** Target genes of CAMTAs and CBF2 detected by ChIP-seq. Blue tick marks indicate genes at which a 6-fold increase in occupancy is observed for CAMTAs during 1-hour treatments with 30 μM CHX at 22°C or 0.1% DMSO at 4°C relative to treatments with 0.1% DMSO at 22°C (mock control) or for CBF2 relative to the wholegenome average during a 4-hour incubation at 4°C. Red-green shading indicates log2 fold-changes in gene expression after 2 hours with the above CHX or cold treatments relative to mock controls, for clusters of genes from *Figure 2.*

**Supplementary Figure 4:**
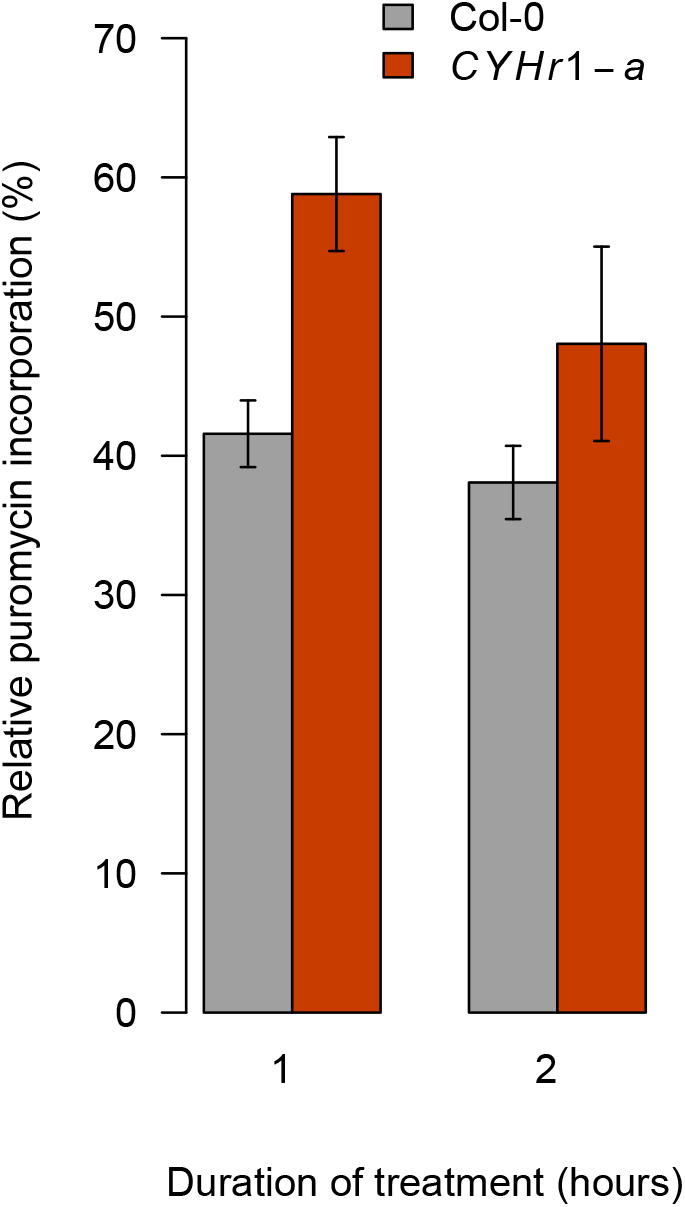
*CYHr1* seedlings have higher translation rates than Col-0 in the presence of CHX. Col-0 and *CYHr1* seedlings (line *CYHr1-a)* were grown in long days at 22°C for 7 days and treated with 30 μM CHX or 0.1% DMSO (mock) at 22°C for 1 or 2 hours, followed by a 30-minute treatment with 150 μM puromycin. The amount of puromycin-labelled proteins was normalised to actin levels and is given for CHX-treated samples relative to mock controls for each time-point. Error bars indicate the standard deviation for 3 biological replicates, with 10 to 15 seedlings per replicate.

**Supplementary Figure 5:**
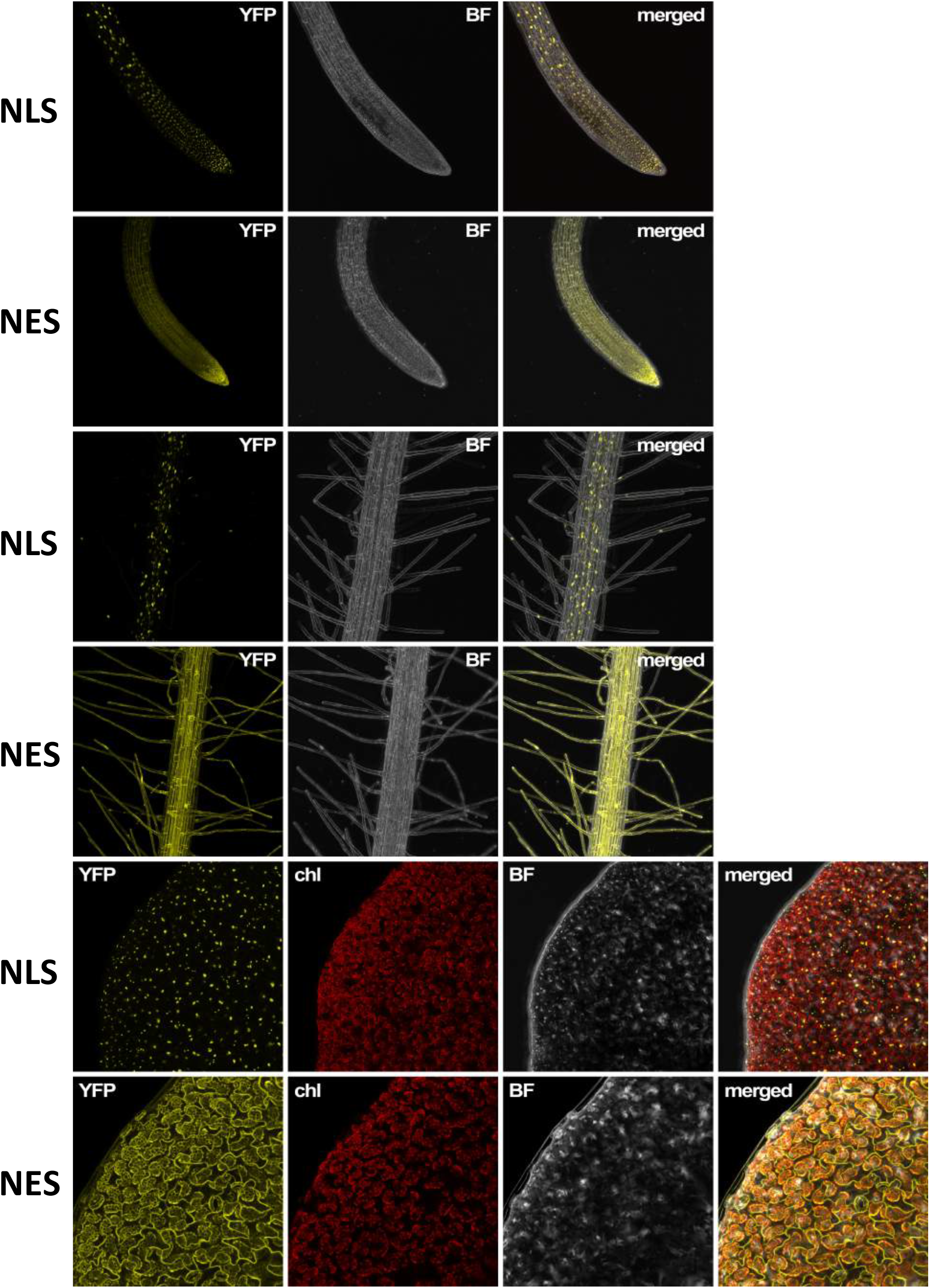
Validation of nuclear- and cytosolic-localised aequorin reporters. Confocal images of the fluorescent protein Venus (modified yellow fluorescent protein, YFP) in *A. thaliana* Col-0 *pUBQ10::NLS_SV40_::VENUS::APOAEQUORIN* (NLS) and *pUBQ10::NES_PK1α_::VENUS::APOAEQUORIN* (NES) roots and cotyledons. Seedlings were grown on agar in long days at 22°C for 7 days before imaging. BF: bright field; chl: chlorophyll autofluorescence.

**Supplementary Figure 6:**
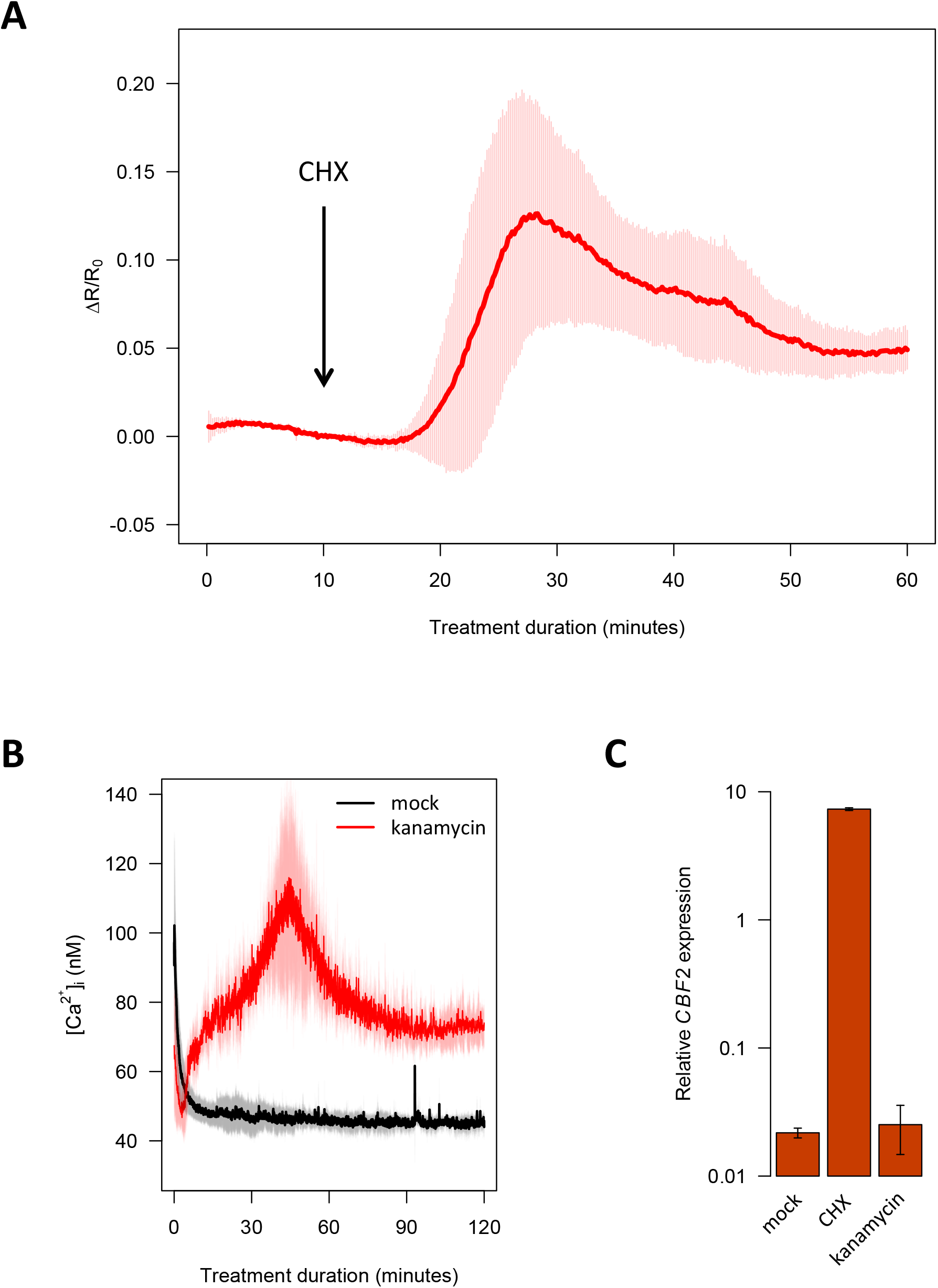
The CHX-induced increase in cytosolic free calcium levels specifically activates early cold signalling. **A:** CHX induces an increase in cytosolic free calcium, as observed using the FRET-based calcium sensor Yellow Cameleon *3.6. A. thaliana* Col-0 *pUBQ10-NES::YC3.6* seedlings were grown on halfstrength MS medium at 22°C with a 16-hour long-day photoperiod for 7 days and placed in dedicated chambers overlaid with cotton wool soaked in imaging solution. The treatment was carried out by supplementing the imaging solution with CHX to a final concentration of 30 μM after 10 minutes (indicated with an arrow). The FRET cpVenus/CFP ratio (R) was measured over time in root tip meristematic cells and normalised to the initial ratio (R_0_). Relative cytosolic free calcium levels, represented by ΔR/R_0_ values, are given for 3 individual seedlings, with error bars indicating standard deviations. **B:** The 70S translation inhibitor kanamycin triggers an increase in intracellular calcium levels. *A. thaliana* Col-0 *pCaMV35S::APOAEQUORIN* seedlings were grown in liquid culture at 20°C with a 12-hour photoperiod for 8 to 12 days and intracellular free calcium levels were quantified luminometrically during 2-hour treatments with 30 μM kanamycin or 0.1% DMSO (mock). Cuvettes were placed in the luminometer immediately after the addition of chemicals. Shading indicates the standard deviation for at least 3 biological replicates, each comprising a cuvette with 3 seedlings. **C:** Kanamycin does not induce *CBF2* expression. *A. thaliana* Col-0 seedlings were grown in liquid culture in long days at 22°C for 7 days and treated with 30 μM kanamycin, 30 μM CHX or 0.1% DMSO (mock) at 22°C for 2 hours. *CBF2* expression was measured by quantitative PCR and normalised to transcript levels of *PP2A* and *UBC21.* Error bars indicate the standard deviation for 3 biological replicates, with 10 to 15 seedlings per replicate.

## Supplementary Movie

**Supplementary Movie:**
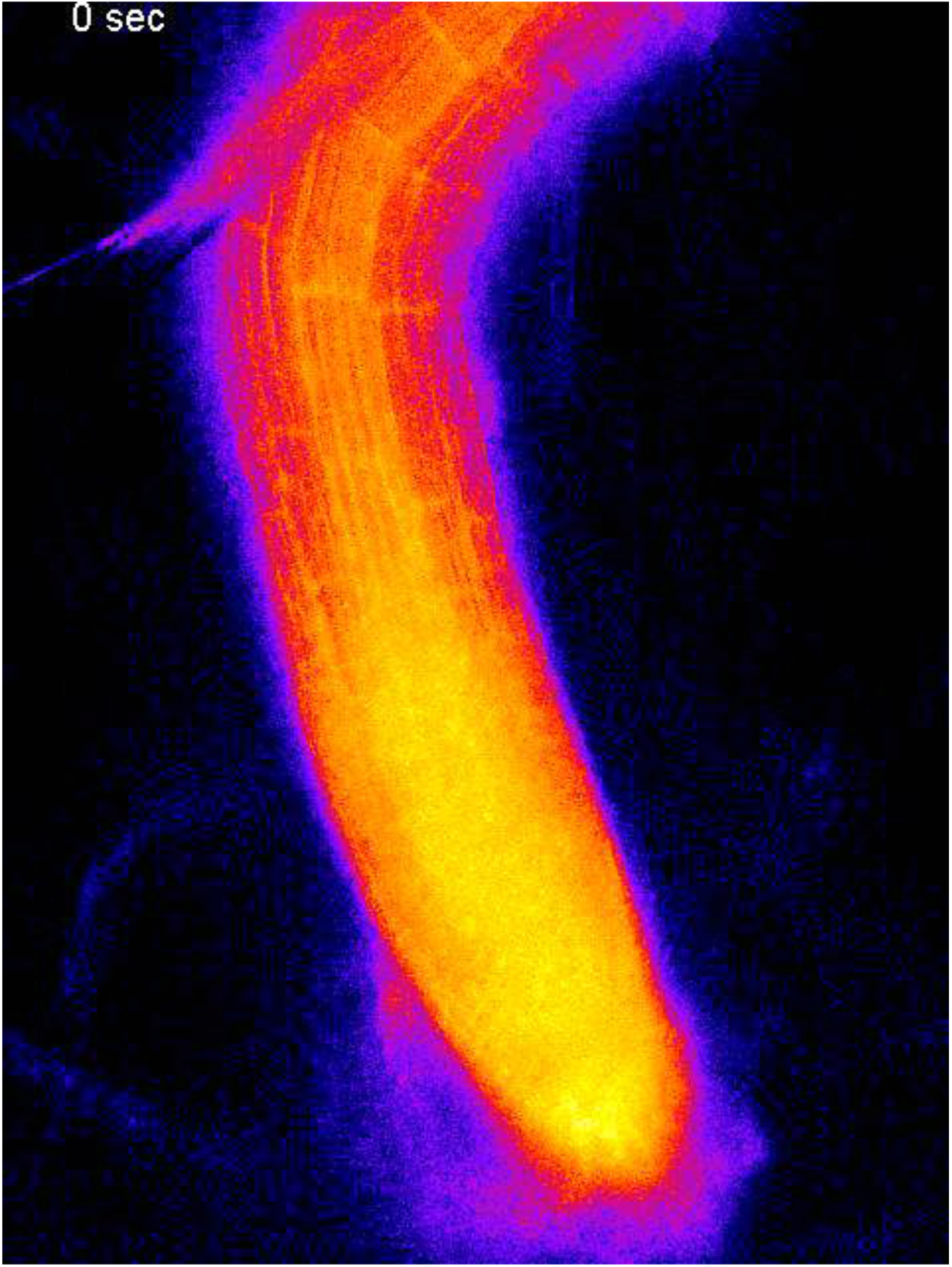
Ratiometric FRET cpVenus/CFP false-colour (LUT: Fire) movie of a representative *A. thaliana* Col-0 *pUBQ10-NES::YC3.6* seedling root tip during treatment with 30 μM CHX treatment, from *Supp. Fig. 6A.*

## References and Notes

Berberich, T. and Kusano, T. (1997). Cycloheximide induces a subset of low temperature-inducible genes in maize. Mol. Gen. Genet. 254: 275–83.

Bouché, N., Yellin, A., Snedden, W.A., and Fromm, H. (2005). PLANT-SPECIFIC CALMODULIN-BINDING PROTEINS. Annu. Rev. Plant Biol. 56: 435–466.

Doherty, C.J., Van Buskirk, H.A., Myers, S.J., and Thomashow, M.F. (2009). Roles for Arabidopsis CAMTA transcription factors in cold-regulated gene expression and freezing tolerance. Plant Cell 21: 972–84.

EK, S., G, C., M, C., and P, P. (2009). SUnSET, a Nonradioactive Method to Monitor Protein Synthesis. Nat. Methods 6.

Farewell, A. and Neidhardt, F.C. (1998). Effect of temperature on in vivo protein synthetic capacity in Escherichia coli. J. Bacteriol.

Jaglo-Ottosen, K.R., Gilmour, S.J., Zarka, D.G., Schabenberger, O., and Thomashow, M.F. (1998). Arabidopsis CBF1 overexpression induces COR genes and enhances freezing tolerance. Science (80-.). 280: 104–106.

Kawai, S., Murao, S., Mochizuki, M., Shibuya, I., Yano, K., and Takagi, M. (1992). Drastic alteration of cycloheximide sensitivity by substitution of one amino acid in the L41 ribosomal protein of yeasts. J. Bacteriol. 174: 254–262.

Kidokoro, S., Maruyama, K., Nakashima, K., Imura, Y., Narusaka, Y., Shinwari, Z.K., Osakabe, Y., Fujita, Y., Mizoi, J., Shinozaki, K., and Yamaguchi-Shinozaki, K. (2009). The Phytochrome-Interacting Factor PIF7 Negatively Regulates DREB1 Expression under Circadian Control in Arabidopsis. Plant Physiol. 151: 2046–2057.

Kidokoro, S., Yoneda, K., Takasaki, H., Takahashi, F., Shinozaki, K., and Yamaguchi-Shinozaki, K. (2017). Different Cold-Signaling Pathways Function in the Responses to Rapid and Gradual Decreases in Temperature. Plant Cell 29: 760–774.

Kim, Y., Park, S., Gilmour, S.J., and Thomashow, M.F. (2013). Roles of CAMTA transcription factors and salicylic acid in configuring the low-temperature transcriptome and freezing tolerance of Arabidopsis. Plant J. 75: 364–376.

Knight, H., Trewavas, A.J., and Knight, M.R. (1996). Cold calcium signaling in Arabidopsis involves two cellular pools and a change in calcium signature after acclimation. Plant Cell 8: 489–503.

Lee, C. and Thomashow, M.F. (2012). Photoperiodic regulation of the C-repeat binding factor (CBF) cold acclimation pathway and freezing tolerance in Arabidopsis thaliana.: 6–11.

Monroy, A.F., Sarhan, F., and Dhindsa, R.S. (1993). Cold-Induced Changes in Freezing Tolerance, Protein Phosphorylation, and Gene Expression (Evidence for a Role of Calcium). Plant Physiol. 102: 1227–1235.

Ohama, N., Kusakabe, K., Mizoi, J., Zhao, H., Kidokoro, S., Koizumi, S., Takahashi, F., Ishida, T., Yanagisawa, S., Shinozaki, K., and Yamaguchi-Shinozaki, K. (2015). The transcriptional cascade in the heat stress response of Arabidopsis is strictly regulated at the expression levels of transcription factors. Plant Cell 28: 181–201.

Park, S., Lee, C.-M., Doherty, C.J., Gilmour, S.J., Kim, Y.. and Thomashow, M.F. (2015). Regulation of the Arabidopsis CBF regulon by a complex low-temperature regulatory network. Plant J. 82: 193–207.

Plieth, C., Hansen, U.-P., Knight, H., and Knight, M.R. (1999). Temperature sensing by plants: the primary characteristics of signal perception and calcium response. Plant J. 18: 491–497.

Polisensky, D.H. and Braam, J. (1996). Cold-shock regulation of the Arabidopsis TCH genes and the effects of modulating intracellular calcium levels. Plant Physiol. 111: 1271–9.

Santagata, S. et al. (2013). Tight coordination of protein translation and HSF1 activation supports the anabolic malignant state. Science 341: 1238303.

Tähtiharju, S., Sangwan, V., Monroy, A.F., Dhindsa, R.S., and Borg, M. (1997). The induction of kin genes in cold-acclimating Arabidopsis thaliana. Evidence of a role for calcium. Planta 203: 442–447.

Yamazaki, T., Kawamura, Y., Minami, A., and Uemura, M. (2008). Calcium-Dependent Freezing Tolerance in Arabidopsis Involves Membrane Resealing via Synaptotagmin SYT1. Plant Cell 20: 3389–3404.

Zarka, D.G., Vogel, J.T., Cook, D., and Thomashow, M.F. (2003). Cold induction of Arabidopsis CBF genes involves multiple ICE (inducer of CBF expression) promoter elements and a cold-regulatory circuit that is desensitized by low temperature. Plant Physiol. 133: 910–8.

## Supplementary references

Kim, Y., Park, S., Gilmour, S. J., & Thomashow, M. F. (2013). Roles of CAMTA transcription factors and salicylic acid in configuring the low-temperature transcriptome and freezing tolerance of Arabidopsis. The Plant Journal, 75(3), 364–376. https://doi.org/10.1111/tpj.12205

Kidokoro, S., Yoneda, K., Takasaki, H., Takahashi, F., Shinozaki, K., & Yamaguchi-Shinozaki, K. (2017). Different Cold-Signaling Pathways Function in the Responses to Rapid and Gradual Decreases in Temperature. The Plant Cell, 29(4), 760–774. https://doi.org/10.1105/TPC.16.00669

Nakamichi, N., Kita, M., Ito, S., Yamashino, T., & Mizuno, T. (2005). PSEUDO-RESPONSE REGULATORS, PRR9, PRR7 and PRR5, Together play essential roles close to the circadian clock of Arabidopsis thaliana. Plant and Cell Physiology, 46(5), 686–698. https://doi.org/10.1093/pcp/pci086

Xu, X., Hotta, C. T., Dodd, A. N., Love, J., Sharrock, R., Young, W. L., Xie, Q., Johnson, C. H., & Webb, A. A. R. (2007). Distinct light and clock modulation of cytosolic free Ca2+ oscillations and rhythmic CHLOROPHYLL A/B BINDING PROTEIN2 promoter activity in Arabidopsis. Plant Cell, 19(11), 3474–3490. https://doi.org/10.1105/tpc.106.046011

Krebs, M., Held, K., Binder, A., Hashimoto, K., Den Herder, G., Parniske, M., Kudla, J., & Schumacher, K. (2012). FRET-based genetically encoded sensors allow high-resolution live cell imaging of Ca 2+ dynamics. Plant Journal, 69(1), 181–192. https://doi.org/10.1111/j.1365-313X.2011.04780.x

Clough, S. J., & Bent, A. F. (1998). Floral dip: A simplified method for Agrobacterium-mediated transformation of Arabidopsis thaliana. Plant Journal, 16(6), 735–743. https://doi.org/10.1046/j.1365-313X.1998.00343.x

Schmidt, E. K., Clavarino, G., Ceppi, M., & Pierre, P. (2009). SUnSET, a nonradioactive method to monitor protein synthesis. Nature Methods, 6(4), 275–277. https://doi.org/10.1038/nmeth.1314

Li, C., & Evans, R. M. (1997). Ligation independent cloning irrespective of restriction site compatibility. Nucleic Acids Research, 25(20), 4165–4166. http://www.ncbi.nlm.nih.gov/pubmed/9321675

Karimi, M., De Meyer, B., & Hilson, P. (2005). Modular cloning in plant cells. Trends in Plant Science, 10(3), 103–105. https://doi.org/10.1016/j.tplants.2005.01.008

Mehlmer, N., Parvin, N., Hurst, C. H., Knight, M. R., Teige, M., & Vothknecht, U. C. (2012). A toolset of aequorin expression vectors for in planta studies of subcellular calcium concentrations in Arabidopsis thaliana. Journal of Experimental Botany, 63(4), 1751–1761. https://doi.org/10.1093/jxb/err406

Box, M. S., Coustham, V., Dean, C., & Mylne, J. S. (2011). Protocol: A simple phenol-based method for 96-well extraction of high quality RNA from Arabidopsis. Plant Methods, 7(1), 1–10. https://doi.org/10.1186/1746-4811-7-7

Clontech Laboratories (2009) Preparation of Yeast Protein Extracts. In: Yeast Protocols Handbook. Protocol No. PT3024-1. Mountain View, CA: Takara Bio. Ch.4.

Pasternak, T., Tietz, O., Rapp, K., Begheldo, M., Nitschke, R., Ruperti, B., & Palme, K. (2015). Protocol: An improved and universal procedure for whole-mount immunolocalization in plants. Plant Methods, 11 (1), 1–10. https://doi.org/10.1186/s13007-015-0094-2

Fricker, M. D., Plieth, C., Knight, H., Blancaflor, E., Knight, M. R., White, N. S. & Gilroy, S. (1999) Fluorescence and Luminescence Techniques to Probe Ion Activities in Living Plant Cells. In: Mason, W.T., ed. (1999) Fluorescent and Luminescent Probes for Biological Activity. San Diego, USA: Academic Press.

Behera, S., Xu, Z., Luoni, L., Bonza, M. C., Doccula, F. G., De Michelis, M. I., Morris, R. J., Schwarzländer, M., & Costa, A. (2018). Cellular Ca 2+ signals generate defined pH signatures in plants. Plant Cell, 30(11), 2704–2719. https://doi.org/10.1105/tpc.18.00655

Jaeger KE, Pullen N, Lamzin S, Morris RJ, Wigge PA (2013). Interlocking feedback loops govern the dynamic behavior of the floral transition in Arabidopsis. Plant Cell. 2013 Mar;25(3):820–33. doi: 10.1105/tpc.113.109355. Epub 2013 Mar 29.

